# ARHGEF6-dependent cytoskeletal regulation underlies a conserved program of forebrain interneuron development

**DOI:** 10.64898/2026.03.09.710568

**Authors:** Carla Liaci, Beatrice Savarese, Elena Ferretti, Jean-Paul Urenda, Junyu Joanna Lu, Giovanni Catapano, Lucia Prandi, Mattia Camera, Rohin Manohar, Simona Rando, Alessandro Umbach, Enis Hidisoglu, Giuseppe Chiantia, Andrea Marcantoni, Maurizio Giustetto, Roberto Oleari, Alyssa Paganoni, Anna Cariboni, Van Truong, Luciano Conti, Giorgia Quadrato, Giorgio R. Merlo

## Abstract

The molecular programs coordinating inhibitory interneuron migration, maturation, and survival during forebrain development remain incompletely understood. Here we investigate ARHGEF6, a RAC1/CDC42 guanine nucleotide exchange factor linked to X-linked intellectual disability (XLID46) and previously studied only at postsynaptic compartments, and reveal an earlier, conserved role in forebrain interneuron development. ARHGEF6 is selectively enriched in the inhibitory lineage during the peak of interneuron generation and migration. Its loss in mice reduces the number of cortical and hippocampal interneurons, disrupts tangential migration, increases developmental cell death, and impairs morphological and electrophysiological maturation. Strikingly, *ARHGEF6*-knockout human iPSC-derived organoids and assembloids mirror these deficits exhibiting increased apoptosis, reduced neuronal output, disorganized growth cones, impaired neurite branching, and disrupted migratory dynamics. These cross-species findings reframe ARHGEF6 as an early, essential orchestrator of inhibitory circuit assembly and reveal a conserved cytoskeletal program whose disruption produces the excitatory-inhibitory imbalance linked to cognitive dysfunction.

## Introduction

The assembly of functional forebrain circuits depends on the precise coordination of three linked processes during development: the directed migration of newborn neurons to their target destinations, the elaboration of their dendritic and axonal arbors, and their survival through developmental checkpoints governed by activity and connectivity (Southwell et al., 2012; Heck et al., 2008; Wong et al., 2018; Causeret et al., 2018; Wong & Marín, 2019). All three processes require dynamic remodeling of the actin cytoskeleton, and all three are disproportionately sensitive to disruption in GABAergic interneurons (INs), which must travel extraordinary distances from their birthplace in the ventral telencephalon to populate the developing cortex and hippocampus (Marin & Rubenstein, 2001; Wonders & Anderson, 2006; Marín, 2013). Despite representing only a minority of the total neuronal population, cortical and hippocampal INs regulate virtually all aspects of local circuit maturation and activity through their morphological, neurochemical, and functional diversity (Pelkey et al., 2017; Lim et al., 2018).

The Rho family of small GTPases, including RHOA, RAC1, and CDC42, are the principal molecular switches that couple extracellular guidance signals to cytoskeletal rearrangements during cell migration and morphogenesis. They cycle between an inactive GDP-bound state and an active GTP-bound state under tight control by guanine nucleotide exchange factors (GEFs), which promote activation, GTPase-activating proteins (GAPs), which stimulate GTP hydrolysis and inactivation, and guanine nucleotide dissociation inhibitors (GDIs), which stabilize the inactive GDP-bound state and prevent reactivation. In neurons, this regulatory network controls axon extension and guidance, growth cone formation and rearrangement, dendritic arborization, and the structural and functional plasticity of synapses (Govek et al., 2011; Tejada-Simon, 2015; Martino et al., 2013; Liaci et al., 2021). Importantly, RAC1 and its regulators play early roles in telencephalic progenitors, influencing self-renewal and cell fate decisions, including the transition of inhibitory progenitors from G1 to S phase via cyclin D and Rb phosphorylation (Vidaki et al., 2012; Lian et al., 2019; Hass et al., 2025). Notably, hundreds of genes have been associated with intellectual disability (ID) (Gupta, 2023; Maia et al., 2021), many of which, including *PAK3*, *ARHGEF6*, *ARHGEF7*, *ARHGEF9*, *OPHN1*, *TRIO*, and *FGD1*, encode regulators and effectors of the Rho family of small GTPases (Liaci et al., 2021).

ARHGEF6 (α-PIX/cool-2) is a guanine nucleotide exchange factor for RAC1 and CDC42 that is well-positioned to coordinate this cytoskeletal program. It contains a tandem Dbl homology (DH)–pleckstrin homology (PH) domain pair conferring catalytic GEF activity, an SH3 domain that recruits p21-activated kinases PAK1–3, and it associates with the ARF GAP scaffold proteins GIT1 and GIT2, placing ARHGEF6 at the intersection of RAC1/CDC42 and ARF GTPase signaling (Manser et al., 1998; Zhou et al., 2016). In the mouse hippocampus, ARHGEF6 localizes within dendritic spines and colocalizes with the postsynaptic marker PSD95, consistent with association with postsynaptic density complexes; its loss or depletion alters spine morphology and density, as well as synaptic plasticity, including reduced long-term potentiation and enhanced long-term depression at the Schaffer collateral–CA1 synapse (Nodé-Langlois et al., 2006; Ramakers et al., 2012). ARHGEF6 has also been proposed to participate in Reelin-dependent Golgi positioning during apical dendrite selection (Meseke et al., 2013). These studies established a postsynaptic role for ARHGEF6 in hippocampal pyramidal neurons, but its function during earlier stages of forebrain development, and specifically in the inhibitory lineage, had not been explored.

Variants in ARHGEF6 have been proposed as the genetic basis for X-linked intellectual disability type 46 (XLID46/MRX46; OMIM 300436), initially identified through a reciprocal X;21 translocation and a segregating intronic variant in a large Dutch pedigree (Kutsche et al., 2000; Yntema et al., 1998). However, subsequent population-level re-evaluation found the reported intronic variant at appreciable frequency in hemizygous males, and clinical curation efforts have accordingly questioned the strength of evidence for a direct causal relationship (Genomics England PanelApp; Piton et al., 2013). This genetic uncertainty is not unusual for rare X-linked conditions, where definitive human genetic evidence may remain elusive for years. What has been missing is direct functional evidence in human neurons. Crucially, the broader RAC1 regulatory network is robustly implicated in neurodevelopmental disorders: variants in RAC1 itself cause a syndrome with intellectual disability, cortical malformations, and developmental delay (Reijnders et al., 2017; Banka et al., 2022), reinforcing the notion that disruption of this signaling axis has major developmental consequences in the human brain.

Here, we show that *ARHGEF6* expression is enriched in the GABAergic inhibitory lineage of the developing mouse and human telencephalon, revealing a developmental pattern distinct from its previously described postsynaptic role. We demonstrate that ARHGEF6 is required for the tangential migration, morphological maturation, electrophysiological development, and survival of forebrain INs in the mouse, and that these requirements are conserved in human ventral forebrain organoids and dorsal–ventral assembloids derived from *ARHGEF6*-knockout (KO) human induced pluripotent stem cells. Together, our findings identify ARHGEF6 as a conserved regulator of IN development and provide a cellular framework for how its disruption may impair inhibitory circuit formation and contribute to cognitive dysfunction. More broadly, this study highlights the value of cross-species models for defining developmental mechanisms relevant to neurodevelopmental disease.

## Results

### 1. Developmental and cell type-specific expression of *ARHGEF6* in the telencephalon

To initially investigate the potential developmental impact of *ARHGEF6* in the human brain, particularly in light of its implication in the pathoetiology of ID, we analyzed publicly available transcriptomic datasets. Bulk RNA-sequencing data from human fetal brain samples at post-conception weeks (PCW) 8–9 revealed *ARHGEF6* is expressed across multiple brain regions (Allen Human Brain Atlas: BrainSpan, Atlas of the Developing Brain; RRID:SCR_008083); **Fig. 1A**). Expression levels were relatively elevated in ventral telencephalic regions, including the ganglionic eminences (GEs), particularly the medial (MGE) and caudal (CGE) ganglionic eminences (**Fig. 1A**), which serve as the principal sources of cortical GABAergic INs during early neurodevelopment. This developmental stage is characterized by intense proliferative activity within the GEs, the onset of IN production, and the early phases of their tangential migration toward the developing cortex (Hansen et al., 2013; Paredes et al., 2016; Keefe et al., 2023). Elevated expression was also observed in the upper rhombic lip (URL) and in the amygdala (AMY), consistent with the developmental origin of several of its nuclei, such as the central and medial amygdala, which are comprised of GE-derived progenitor domains that predominantly generate GABAergic neurons (Aerts et al., 2021; Prakash et al., 2025).

**Figure 1.**
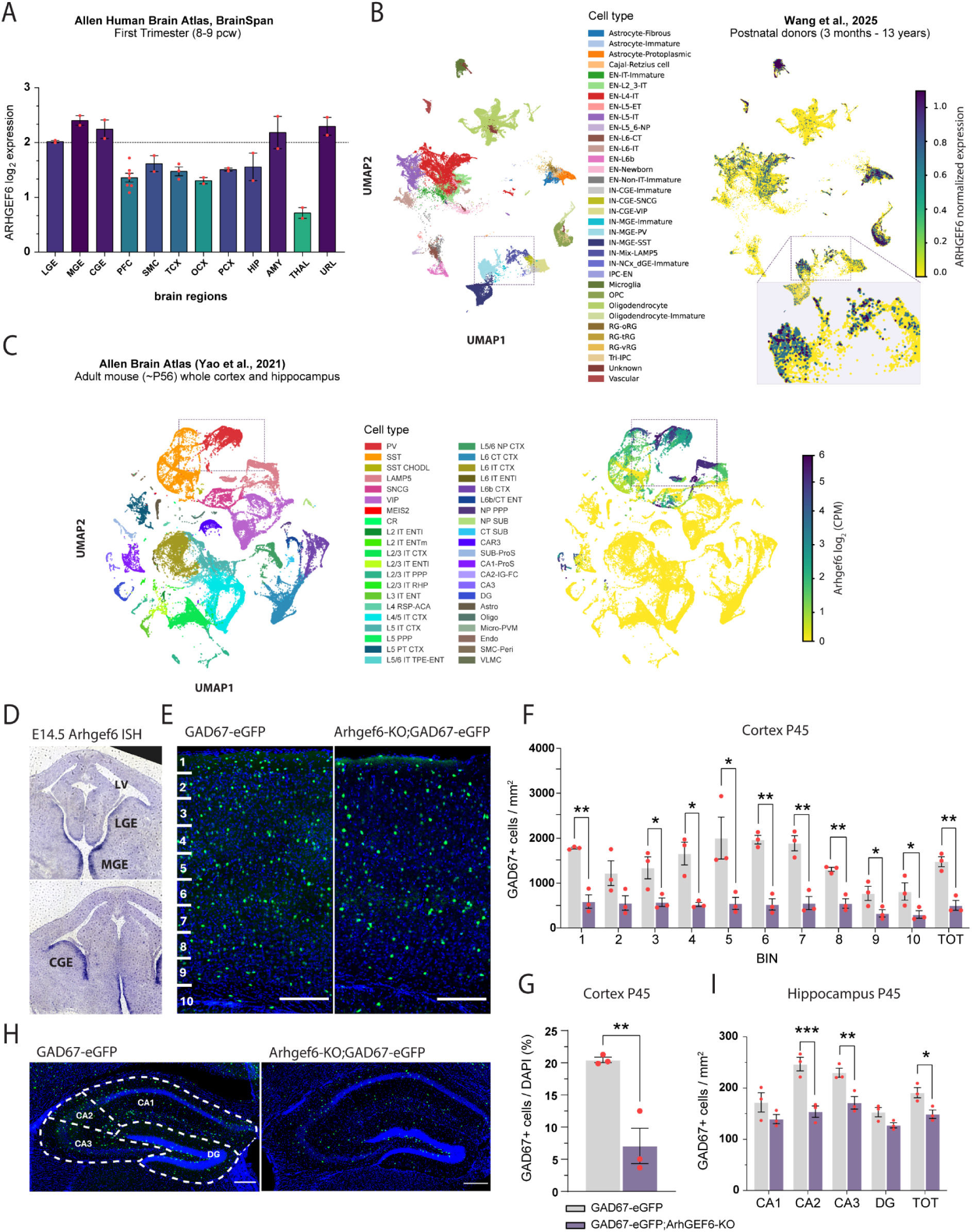
*ARHGEF6* expression in the ventral telencephalic lineage and reduced GABAergic IN number in the *Arhgef6*-*KO* murine telencephalon. **A.** *ARHGEF6* expression across brain regions between 8-9 post-conception weeks from bulk RNA-seq data. 30 total samples from 2 unique donors. The dashed line indicates the minimum level of expression detected in the ganglionic eminences (GEs). PFC, prefrontal cortex (areas: dorsolateral prefrontal cortex, anterior [rostral] cingulate [medial prefrontal] cortex, orbital frontal cortex, ventrolateral prefrontal cortex); SMC, sensorimotor cortex (areas: primary motor cortex [area M1, area 4], primary somatosensory cortex [area S1, areas 3, 1, 2], primary motor-sensory cortex); TCX, temporal cortex (areas: temporal neocortex, posterior [caudal] superior temporal cortex, inferolateral temporal cortex [area TEv, area 20], primary auditory cortex [core]); OCX, occipital cortex (areas: occipital neocortex, primary visual cortex [striate cortex, area V1/17]); PCX, parietal cortex (areas: parietal neocortex, posteroventral [inferior] parietal cortex); HIP, hippocampus (areas: hippocampus [hippocampal formation]); AMY, amygdala (areas: amygdaloid complex); STR, striatum (areas: striatum); THAL, thalamus (areas: dorsal thalamus, mediodorsal nucleus of thalamus); CBL/URL, cerebellum, upper rhombic lip (areas: cerebellum, cerebellar cortex, upper [rostral] rhombic lip). **B.** UMAP of single-nucleus RNA-sequencing (snRNA-seq) data from human postnatal donors (3 months - 13 years) colored by cell type. UMAP of snRNA-seq data with each nuclei’s *ARHGEF6* expression. Expression values were clipped at the 99th percentile to improve color scale contrast and reduce the influence of outlier cells. 114,216 cells included from 18 samples across 9 unique subjects. EN-IT-Immature, intratelencephalic excitatory immature neurons; EN-L2_3-IT, layer 2/3 intratelencephalic excitatory neurons; EN-L4-IT, layer 4 intratelencephalic excitatory neurons; EN-L5-ET, layer 5 extratelencephalic excitatory neurons; EN-L5-IT, layer 5 intratelencephalic excitatory neurons; EN-L5_6-NP, layer 5/6 near-projecting excitatory neurons; EN-L6-CT, layer 6 corticothalamic excitatory neurons; EN-L6-IT, layer 6 intratelencephalic excitatory neurons; EN-L6b, layer 6b excitatory neurons; EN-Newborn, Newborn excitatory neurons; EN-Non-IT-Immature, immature non-intratelencephalic excitatory neuron;. IN-CGE-Immature, immature caudal ganglionic eminence-derived inhibitory neurons; IN-CGE-SNCG, immature caudal ganglionic eminence-derived gamma-synuclein inhibitory neurons; IN-CGE-VIP, caudal ganglionic eminence-derived vasoactive intestinal polypeptide inhibitory neurons; IN-MGE-Immature, immature medial ganglionic eminence-derived inhibitory neurons; IN-MGE-PV, medial ganglionic eminence-derived parvalbumin inhibitory neurons; IN-MGE-SST, medial ganglionic eminence-derived somatostatin inhibitory neurons; IN-Mix-LAMP5, mixed lysosomal-associated membrane protein family member 5 inhibitory neurons; IN-NCx_dGE-Immature, immature neocortex and dorsal ganglionic eminence-derived inhibitory neurons; IPC-EN, intermediate progenitor cell for excitatory neurons; OPC, oligodendrocyte precursor cells; RG-oRG, outer radial glial cells; RG-tRG, truncated radial glial cells; RG-vRG, ventricular radial glial cells; Tri-IPC, tripotential intermediate progenitor cells. **C**. Representative *in situ* hybridization (ISH) images using an *Arhgef6* antisense RNA probe on coronal sections of embryonic (E) 14.5 wild-type (WT) mouse brains. LV, lateral ventricle; LGE, lateral ganglionic eminence; MGE, medial ganglionic eminence; CGE, caudal ganglionic eminence. Scale bar: 400 µm. **D**. UMAP of single-cell RNA-sequencing (scRNA-seq) data from ∼8 week old (∼P56) mice colored by cell type. UMAP of scRNA-seq data with *Arhgef6* expression for each cell cluster. 73347 cells included. PV, parvalbumin; SST, somatostatin; SST CHOLD, somatostatin and chondrolectin; LAMP5, lysosomal-associated membrane protein family member 5; SNCG, gamma-synuclein; VIP, vasoactive intestinal polypeptide; MEIS2, meis homeobox 2; CR, cajal-retzius cell; L2 IT ENTl, layer 2 intratelencephalic lateral entorhinal area; L2 IT ENTm, layer 2 intratelencephalic medial entorhinal area; L2/3 IT CTX, layer 2/3 intratelencephalic isocortex; L2/3 IT ENTl, layer 2/3 intratelencephalic lateral entorhinal area; L2/3 IT PPP, layer 2/3 intratelencephalic postsubiculum-presubiculum-parasubiculum; L2/3 IT RHP, layer 2/3 intratelencephalic retrohippocampal region; L3 IT ENT, layer 3 intratelencephalic entorhinal area; L4 RSP-ACA, layer 4 retrosplenial area-anterior cingulate area; L4/5 IT CTX, layer 4/5 intratelencephalic isocortex; L5 IT CTX, layer 5 intratelencephalic isocortex; L5 PPP, layer 5 postsubiculum-presubiculum-parasubiculum; L5 PT CTX, layer 5 pyramidal tract isocortex; L5/6 IT TPE-ENT, layer 5/6 intratelencephalic temporal association areas-perirhinal area-ectorhinal area-entorhinal area; L5/6 NP CTX, layer 5/6 near-projecting isocortex; L6 CT CTX, layer 6 corticothalamic isocortex; L6 IT CTX, layer 6 intratelencephalic isocortex; L6 IT ENTl, layer 6 intratelencephalic lateral entorhinal area; L6b CTX, layer 6b isocortex; L6b/CT ENT, layer 6b corticothalamic entorhinal area; NP PPP, near-projecting postsubiculum-presubiculum-parasubiculum; NP SUB, near-projecting subiculum; CT SUB, corticothalamic subiculum; Car3, carbonic anhydrase 3; SUB-Pros, subiculum-prosubiculum; CA1-ProS, field cornu ammonis 1-prosubiculum; CA2-IG-FC, field cornu ammonis 2-fasciola cinerea-indusium griseum; CA3, field cornu ammonis 3; DG, dentate gyrus; Astro, astrocyte; Oligo, oligodendrocyte; Micro-PVM, microglia/perivascular macrophage; Endo, endothelial cell; SMC-Peri, smooth muscle cell perivascular area; VLMC, vascular leptomeningeal cell. **E**. Representative maximum intensity projection of z-stack images (30 serial image planes, z-step size = 1 µm) of the somatosensory cortex of postnatal (P) 45 *GAD67-eGFP* (left) and *GAD67-eGFP;Arhgef6-KO* mice (right). Scale bars, 200 µm. **F**. Average density of eGFP^+^ GABAergic INs in the adult cortex in each of the 10 bins (shown in E). At least 3 distinct sections distributed along the anteroposterior axis from 3 different mice per genotype were analyzed. p-values (from bin 1 to 10) = (1) 0.004; (2) 0.058; (3) 0.032; (4) 0.012; (5) 0.033; (6) 0.004; (7) 0.005; (8) 0.005; (9) 0.043; (10) 0.044; (TOT) 0.005. **G**. Average percentage of eGFP^+^ GABAergic INs in the adult cortex over the total number of cells. At least 3 distinct sections distributed along the anteroposterior axis from 3 different mice per genotype were analyzed. p-values = 0.008. **H**. Representative maximum intensity projection of z-stack images (30 serial image planes, z-step size = 1 µm) of the hippocampus of P45 *GAD67-eGFP* (left) and *GAD67-eGFP;Arhgef6-KO* (right) mice. Scale bar, 100 µm. **I**. Average density of eGFP^+^ GABAergic INs in the cornu ammonis region 1, 2, and 3 (CA1, CA2, CA3), and dentate gyrus (DG) regions and whole hippocampus. At least 3 distinct sections distributed along the anteroposterior axis from 3 different mice per genotype were analyzed. p-values = 0.097 (CA1); 0.00004 (CA2); 0.005 (CA3); 0.12 (DG); 0.044 (TOT). p-values were calculated using unpaired multiple t-tests corrected for False Discovery Rate (<1%) (F) and Holm-Šídák method (H). * = p< 0.05, ** = p< 0.01, *** = p< 0.001. Data are presented as mean ± SEM. Each dot represents one animal.

To further examine the cellular distribution of *ARHGEF6* expression, we next analyzed postnatal human cortical single-nucleus RNA sequencing (snRNA-seq) datasets spanning 3 months to 13 years of age (Wang et al., 2025)(**Fig. 1B; Tab. S1**). UMAP visualization of annotated cell populations showed that *ARHGEF6* is expressed within both neuronal and non-neuronal clusters (**Fig. 1B**). Within neuronal populations, *ARHGEF6* is enriched in specific GABAergic lineages, including MGE-derived parvalbumin (PV) INs and CGE-derived LAMP5-expressing INs (as shown in the magnified view in **Fig. 1B**). In contrast, excitatory cells exhibited comparatively lower *ARHGEF6* expression (**Fig. 1B; Tab. S1**). The detection of *ARHGEF6* expression in oligodendrocyte progenitor cells (OPCs) is consistent with the developmental origin of early OPC populations from ventral telencephalic progenitor domains, including the GEs (Cai et al., 2024). In addition to neural lineages, *ARHGEF6* is also expressed in immune-related cell types, including monocytes and monocyte progenitors, as similarly reported in broader transcriptomic datasets containing hematopoietic cells (The Human Protein Atlas; Uhlén et al., 2015; Uhlen et al., 2017). As microglia originate from yolk-sac-derived erythro-myeloid progenitors and share transcriptional programs with monocyte/macrophage lineages, the expression of *ARHGEF6* in this cell type is therefore consistent with its broader expression in cells of the myeloid/monocytic lineage.

Consistent with the early expression of *ARHGEF6* observed in the human developing telencephalon (**Fig. 1A**), when we examined *Arhgef6* expression in coronal sections of mouse embryonic brains using *in situ* hybridization (ISH) with a previously validated probe (Lein et al., 2007), the earliest detectable expression was observed around embryonic day (E) 14.5, with higher amounts in MGE and CGE and comparatively lower expression in the lateral ganglionic eminence (LGE) (**Fig. 1C**). The enrichment of *ARHGEF6* expression in inhibitory lineages was further supported by analysis of the publicly available Allen Brain Atlas single-cell RNA sequencing (scRNA-seq) dataset from adult (∼P56) mouse cortex and hippocampus (Yao et al., 2021). Interestingly, as in the postnatal human brain, enrichment was detected in multiple subtypes of GABAergic INs, particularly MGE-derived PV+ cells and CGE-derived LAMP5+ INs (**Fig. 1D; Tab. S2**). Among excitatory populations, only CA3 pyramidal neurons and mossy cells exhibited expression levels comparable to those observed in inhibitory clusters (**Fig. 1D, Tab. S2**). Elevated expression within the CA3 stratum pyramidale was further confirmed by *in situ* hybridization (ISH) data available through the Allen Brain Atlas portal (Lein et al., 2007; Allen Institute for Brain Science, experiment RP_060315_04_C08). Consistent with these findings, previous studies have reported elevated *Arhgef6* expression in neuropil regions of the adult mouse hippocampus (Ramakers et al., 2012; Meyer, 2014).

Collectively, these findings suggest a previously undefined developmental role for this gene in the GABAergic telencephalic lineage that can be observed across species.

### 2. Arhgef6 deficiency reduces INs numbers in the adult cortex and hippocampus

Given that *ARHGEF6* expression is enriched in GABAergic INs in both the early embryonic and adult brains, we first aimed to determine whether adult mice exhibited any defects. To investigate the effect of *Arhgef6* deficiency on the inhibitory neuronal lineage, *Arhgef6-KO* mice (Ramakers et al., 2012) were crossed with the *GAD67-eGFP* reporter line (Tamamaki et al., 2003).

We then examined coronal sections of the sensory cortex and hippocampus from adult (P45) *GAD67-eGFP;Arhgef6-KO* mice and compared them with sections from *GAD67-eGFP* control mice. The brains of *GAD67-eGFP;Arhgef6*-*KO* mice exhibited a significant reduction in the number of GAD67^+^ cells throughout the cortex (**Fig. 1E-G**), particularly in layers I, V-VI (corresponding to BIN 1, 6-8), and in the hippocampus, specifically within the CA2 and CA3 regions (**Fig. 1H, I**). The decrease in IN number within the CA2–CA3 regions occurs in an area enriched for *Arhgef6* expression (Lein et al., 2007; Allen Institute for Brain Science, experiment RP_060315_04_C08).

These findings suggest that the inhibitory balance may be altered in the adult mouse cortex and hippocampus, complementing previous findings and potentially explaining the cognitive impairment observed in mice (Ramakers et al., 2012).

### 3. Loss of Arhgef6 impairs cell survival during embryonic development

Our observation of a significant reduction in IN numbers in the adult cortex and hippocampus of *Arhgef6-KO* mice suggested that early developmental defects might underlie the adult phenotype. We hypothesized that the reduced number of adult cortical and hippocampal INs may, at least in part, result from impaired cell survival during development. To test this, we performed TUNEL staining on early (E14.5) (**Fig. 2A**) and late (E18.5) (**Fig. 2D**) embryonic brain sections from *Arhgef6-KO* and control mice.

**Figure 2.**
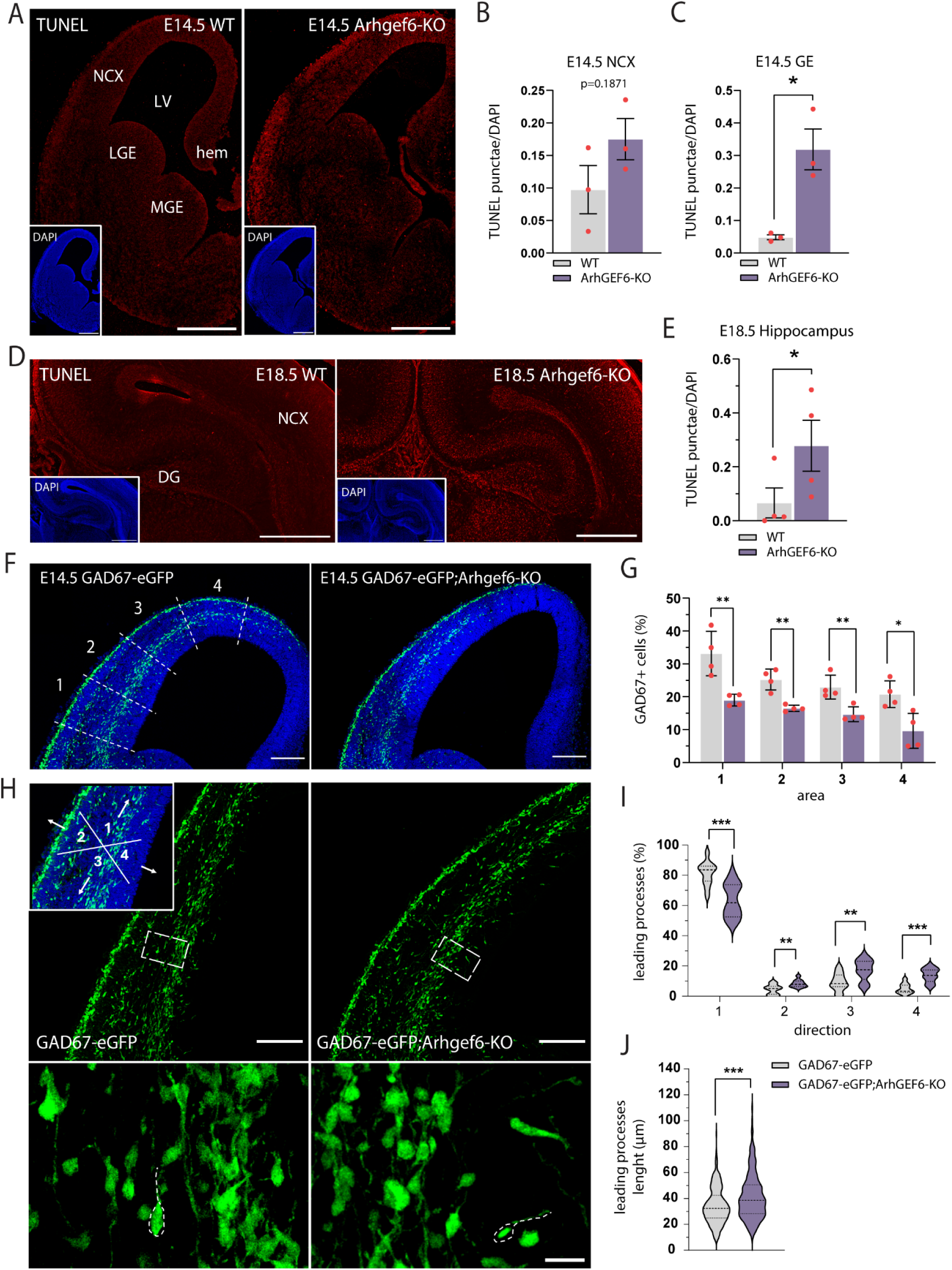
Murine embryonic INs show altered migratory patterns and reduced survival. **A**. Representative maximum intensity projection of z-stack images (30 serial image planes, z-step size = 1 µm) of telencephalic coronal sections from E14.5 WT (left) and *Arhgef6-KO* (right) mouse stained for apoptotic nuclei using the TUNEL assay; insets show the corresponding DAPI staining highlighting overall cytoarchitecture. NCX, neocortex; LV, lateral ventricle; hem, cortical hem; LGE, lateral ganglionic eminence; MGE, medial ganglionic eminence. Scale bars, 500 µm. **B**, **C**. Average percentage of TUNEL punctae over the total number of nuclei. At least 3 distinct sections distributed along the anteroposterior axis from 3 different mice per genotype were analyzed. p-values = 0.1871 (E14.5 NCX); 0.0129 (E14.5 GE). **D**. Representative maximum intensity projection of z-stack images (30 serial image planes, z-step size = 1 µm) of hippocampal coronal sections from E18.5 WT (left) and *Arhgef6-KO* (right) mouse stained for apoptotic nuclei using the TUNEL assay; insets show the corresponding DAPI staining highlighting overall cytoarchitecture. NCX, neocortex; DG, dentate gyrus. Scale bars, 500 µm. **E**. Average percentage of TUNEL punctae over the total number of nuclei. At least 3 distinct sections distributed along the anteroposterior axis from 4 different mice per genotype were analyzed. p-value = 0.029 (E18.5 HIPP). **F**. Representative maximum intensity projection of z-stack images of neocortical coronal sections from E14.5 *GAD67-eGFP* (left) and *GAD67-eGFP;Arhgef6-KO* (right) stained for DAPI. The *GAD67-eGFP* shows the division of the neocortex into four areas of equal size (1, 2, 3, 4) used for the analysis (G). Scale bars, 200 µm. **G**. Average percentage of eGFP^+^ GABAergic INs over the total number of cells in each of the four areas. At least 3 distinct sections distributed along the anteroposterior axis from 4 different mice per genotype were analyzed. p-value (from area 1 to 4) = 0.008 (1); 0.006 (2); 0.008 (3); 0.0119 (4). **H**. Representative maximum intensity projection of z-stack images of neocortices from E14.5 *GAD67-eGFP* (left) and *GAD67-eGFP;Arhgef6-KO* (right) embryos; the inset in the *GAD67-eGFP* shows a schematic representation of the four directions analysed: tangential-dorsal (1), pial (2), ventral (3), ventricular (4). Scale bars, 150 µm. Below, higher magnification of the regions outlined by dashed white boxes, showing different orientations of the leading processes of INs. Scale bar, 50 µm. **I**. Percentage of eGFP^+^ GABAergic INs with the leading processes oriented in each of the 4 directions. p-values (from direction 1 to 4) = 0.0001 (1); 0.002 (2); 0.003 (3); 0.00004 (4). **J**. Average length (µm) of leading processes of migrating eGFP^+^ GABAergic INs. p-value < 0.001. p-values were calculated using unpaired t-test (B, C, E, J), unpaired multiple t-tests corrected for False Discovery Rate (<1%) (G, I). * = p< 0.05, ** = p< 0.01, *** = p< 0.001. Data are presented as mean ± SEM. Each dot represents one animal. Violin plots show the distribution of values with median and quartiles indicated.

At E14.5, TUNEL staining was analyzed in the neocortex and GE (**Fig. 2A–C**). No significant differences were detected in the cortex (**Fig. 2B**). In contrast, a significant increase in TUNEL punctae was observed in the GE of *Arhgef6-KO* embryos compared with controls (**Fig. 2C**), indicating elevated cell death in this region. We next examined apoptosis at a later embryonic (E18.5) timepoint, when the hippocampus is morphologically recognizable and the GE have largely regressed, and found a significant increase in TUNEL punctae in the hippocampus of *Arhgef6-KO* embryos (**Fig. 2D, E**).

These results suggest that loss of Arhgef6 impairs cell survival during embryonic development and may thereby contribute to the postnatal reduction in IN number.

### 4. Arhgef6 loss impairs tangential migration and directionality of GABAergic INs

Given the role of ARHGEF6 as a RAC1 GEF and the requirement of RAC1 signaling for IN migration (Vidaki et al., 2012; Chen et al., 2007; Tivodar et al., 2015), we investigated IN migration in *GAD67-eGFP;Arhgef6-KO* mice.

We examined coronal brain sections from E14.5 *GAD67-eGFP;Arhgef6-KO* embryos and quantified the density of INs undergoing tangential ventral-to-dorsal migration into the neocortex (**Fig. 2F, G**). The neocortex was subdivided into four equal regions, and IN numbers were quantified as previously described (Sun et al., 2021; Eid et al., 2025). We observed a significant reduction in the percentage of INs entering the neocortex in *GAD67-eGFP;Arhgef6-KO* embryos across all four defined regions (1–4) (**Fig. 2G**). We next analyzed the directionality of migrating INs by quantifying the orientation of their leading process relative to the tangential axis (**Fig. 2H, I**). In E14.5 *GAD67-eGFP;Arhgef6-KO* embryos, the proportion of cells oriented dorsally along the main tangential migratory stream (direction 1) was significantly reduced compared with controls (**Fig. 2I**). In contrast, a significant increase was observed in the percentage of INs oriented toward non-tangential directions (directions 2–4), including pial (2), ventral (3), and ventricular (4) orientations (**Fig. 2I**). In the same sections, we measured the length of the leading process in tangentially migrating INs and found a significant increase in *GAD67-eGFP;Arhgef6-KO* embryos compared with controls (**Fig. 2J**).

These findings support an IN migration defect in KO animals, likely reflecting impaired exit from the GE, as previously observed following RAC1 ablation (Chen et al., 2007; Vidaki et al., 2012), and indicate that *Arhgef6* deficiency disrupts leading-process directional coherence while inducing morphological alterations in tangentially migrating cortical INs.

### 5. Arhgef6 deficiency impairs neurite branching in hippocampal INs

Given the well-established role of RAC1 in neurite outgrowth (Liaci et al., 2021), we asked whether reduced RAC1 pathway activity resulting from Arhgef6 loss (Ramakers et al., 2012) affects the morphology of GABAergic INs. To test this, we prepared primary dissociated hippocampal cultures from P0 *GAD67-eGFP* and *GAD67-eGFP;Arhgef6-KO* pups and maintained them for 10 days *in vitro* (DIV) (**Fig. 3A**).

**Figure 3.**
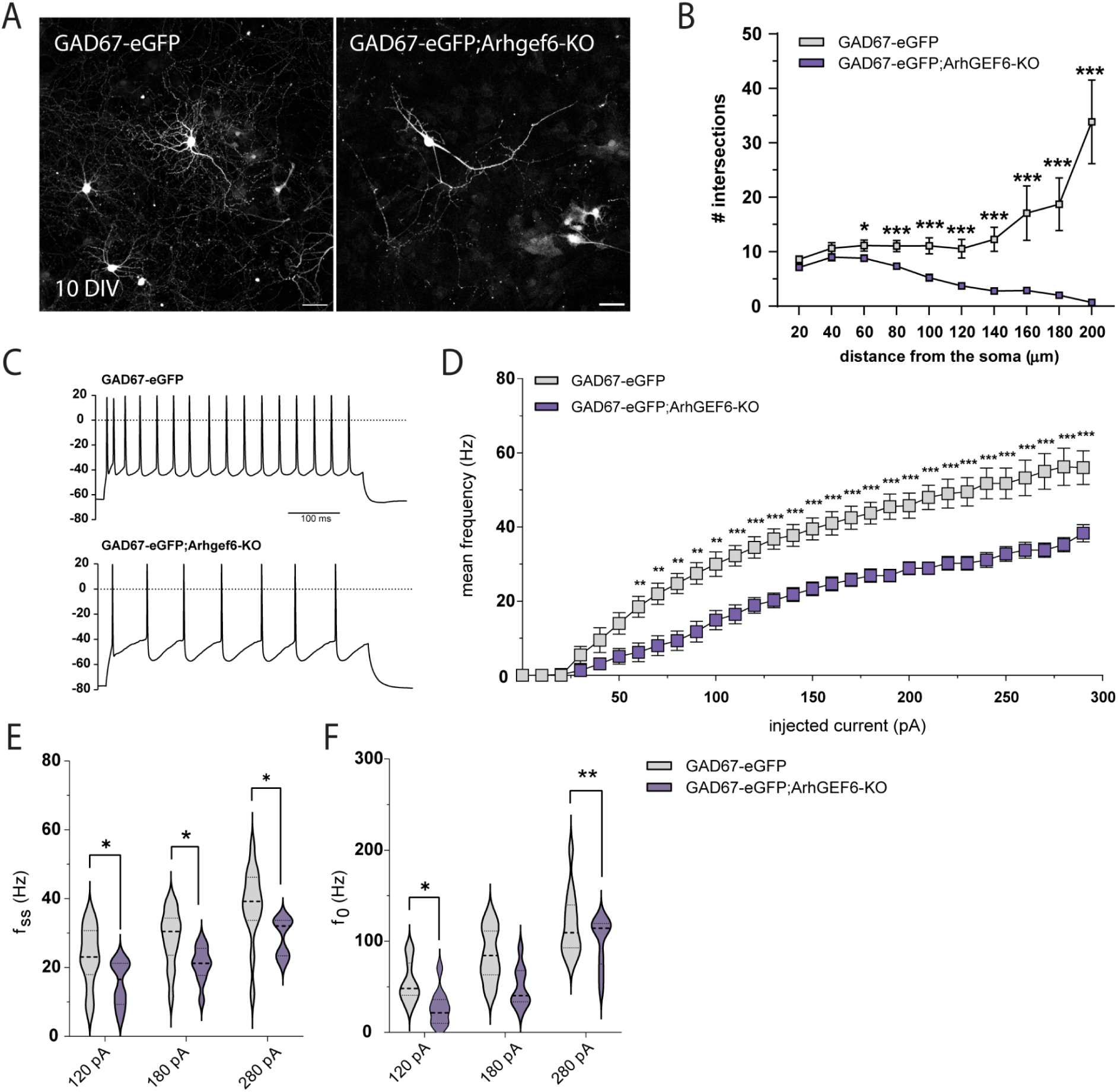
*Arhgef6-KO* INs exhibit simpler branching and reduced excitability. **A**. Representative micrographs of eGFP^+^ primary murine hippocampal INs from *GAD67-eGFP* (left) and *GAD67-eGFP;Arhgef6-KO* (right) after 10 days *in vitro* (DIV). Scale bars, 20 µm. **B**. Sholl analysis showing the overall complexity of arborization in *GAD67-eGFP* and *GAD67-eGFP;Arhgef6-KO* primary INs after 10 DIV. p-values (from 20 to 200 µm) = 0.023 (20 µm), 0.054 (40 µm), 0.019 (60 µm), 0.0009 (80 µm), 0.0003 (100 µm), 0.0003 (120 µm), 0.0003 (140 µm), 0.0009 (160 µm), 0.0007 (180 µm), 0.0003 (200 µm). Around 30 neurons from 2 independent primary cultures were analyzed for each genotype. **C**. Representative whole-cell current clamp recordings of action potentials evoked by 100 pA step current for *GAD67-eGFP* (top) and *GAD67-eGFP;Arhgef6-KO* (bottom) INs. **D**. Average firing frequency vs. current relationships recorded in *GAD67-eGFP* and *GAD67-eGFP;Arhgef6-KO* INs in response to a set of injected current steps (from 0 to 300 pA; 10 pA steps). p-values (from 0 to 300 pA) = 0.999 (0 pA), >0.999 (10 pA), 0.999 (20 pA), 0.708 (30 pA), 0.368 (40 pA), 0.102 (50 pA), 0.008 (60 pA), 0.002 (70 pA), 0.0004 (80 pA), 0.0004 (90 pA), 0.0006 (100 pA), 0.0004 (110 pA), 0.0004 (120 pA), 0.0002 (130 pA), 0.0003 (140 pA), 0.0003 (150 pA), 0.0003 (160 pA), 0.0002 (170 pA), 0.0002 (180 pA), 0.00002 (190 pA), 0.0002 (200 pA), 0.00001 (210 pA), 0.00002 (220 pA), 0.00001 (230 pA), 0.000002 (240 pA), 0.00002 (250 pA), 0.00001 (260 pA), 0.000002 (270 pA), 0.000002 (280 pA), 0.0004 (290 pA). Patch-clamp recordings were obtained from 11 cells from 4 *GAD67-eGFP* mice and 12 cells from 4 *GAD67-eGFP;Arhgef6-KO* mice. **E**. Mean firing frequency at steady state (f_ss_) in response to current injections (120, 180, and 280 pA) in *GAD67-eGFP* and *GAD67-eGFP;Arhgef6-KO* INs. p-values: 0.039 (120 pA), 0.039 (180 pA), 0.024 (280 pA). **F**. Mean initial firing frequency (f_0_) measured at the onset of current injection in *GAD67-eGFP* and *GAD67-eGFP;Arhgef6-KO* INs at 120, 180, and 280 pA. p-values: 0.034 (120 pA), 0.0098 (180 pA), 0.127 (280 pA). p-values were calculated using unpaired multiple t-tests corrected for False Discovery Rate (<1%) (B), and Holm-Šídák method (D, E, F). * = p< 0.05, ** = p< 0.01, *** = p< 0.001. Data are presented as mean ± SEM. Violin plots show the distribution of values with median and quartiles indicated.

At 10 DIV, we assessed neurite length, arborization, and overall complexity using Sholl analysis. Compared with controls, *Arhgef6-KO* GAD67^+^ neurons showed fewer intersections between 80 and 200 μm from the soma, whereas control neurons displayed a progressive increase in intersections with distance, up to 200 μm (**Fig. 3B**). No significant differences were detected in the soma diameter, number of primary neurites, and length of the longest neurite between genotypes (**Fig. S1A-C**).

These results suggest that *Arhgef6* loss compromises distal branching and neurite complexity.

### 6. Arhgef6 loss reduces INs excitability

Previous studies have shown that loss of Arhgef6 in hippocampal pyramidal neurons alters long-term potentiation (LTP) and long-term depression (LTD) in the Schaffer collateral–CA1 pathway (Ramakers et al., 2012). To assess whether Arhgef6 depletion also affects IN electrophysiological properties, we performed whole-cell patch-clamp recordings from GAD67⁺ cells in acute slices from adult *GAD67-eGFP* and *GAD67-eGFP;Arhgef6-KO* mice.

Representative traces of evoked action potentials revealed differences in baseline firing and firing frequency between genotypes (**Fig. 3C**). *Arhgef6*-*KO* INs displayed a reduced firing rate in response to incremental current injections (0–300 pA), with significant differences emerging from 70 pA onward (**Fig. 3D**). We quantified instantaneous firing frequency at spike train onset (f_₀_) and steady state (f_ss_) to evaluate spike frequency adaptation. Both f_₀_ and f_ss_ were significantly reduced at 120 and 280 pA, while only f_ss_ was reduced at 180 pA (**Fig. 3E, F**).

In contrast, input resistance (R_in_), resting membrane voltage (V_rest_), membrane capacitance, and rheobase were unchanged (**Fig. S1D, F, G, H**). Similarly, single action potential properties, including peak amplitude, half-width, maximal rising slope, and maximal repolarizing slope, did not differ between genotypes (**Fig. S1I, J, K, L**).

Overall, these data indicate that Arhgef6 loss reduces IN firing output, consistent with a hypoexcitable phenotype, without overt changes in passive membrane properties or single action potential waveform.

### 7. Establishment of an *ARHGEF6-KO* human iPSC line and cytoskeletal phenotyping in NPCs

To investigate the role of *ARHGEF6* in human development we generated an *ARHGEF6-KO* human induced pluripotent stem cell (hiPSC) line using CRISPR–Cas9.

A commercially available male hiPSC line (ATCC-DYS0100) was targeted with a crRNA designed to disrupt exon 3, the first exon common to all *ARHGEF6* isoforms, resulting in a one–base pair deletion that introduced a frameshift mutation (**Fig. S2A**). As a control, we generated an RNP negative line from the same parental line using a non-targeting crRNA. The *ARHGEF6-KO* line maintained pluripotent stem cell state, as indicated by co-expression of OCT4 and SOX2 (**Fig. S2B**). Sequencing confirmed the absence of mutations at the three top predicted off-target exonic sites in the mutant line (**Fig. S2C**). *ARHGEF6-KO* hiPSC robustly differentiated into embryoid bodies containing derivatives of all three germ layers (ACTA2⁺ mesodermal derivatives, GATA4⁺ endoderm, and MAP2⁺ neuroectoderm), indicating preserved developmental potential (**Fig. S2D**), as well as neural progenitor cells (NPCs) coexpressing NESTIN, SOX2 and the radial glia markers PAX6 and FABP7 (**Fig. S2E**).

To assess the impact of RAC1 pathway disruption on cytoskeletal dynamics, we performed Phalloidin-FITC staining on NPCs to visualize F-actin filaments and evaluate cytoskeletal organization (**Fig. S2F**). We quantified the anisotropy index, which reflects the degree of alignment of F-actin fibers: cells with a greater proportion of fibers oriented along a common axis exhibit higher anisotropy. *ARHGEF6-KO* cells showed a significantly increased anisotropy index (**Fig. S2G**), indicating a more “aligned” actin cytoskeleton than controls. In parallel, we quantified Phalloidin fluorescence to assess total polymerized actin and found that *ARHGEF6-KO* cells displayed a significant reduction in F-actin content compared with controls (**Fig. S2H**).

### 8. ARHGEF6 deficiency impairs ventral forebrain organoid growth and shape

To investigate ARHGEF6 function during human subpallial (ventral telencephalic) development, we used the *ARHGEF6-KO* and RNP negative control hiPSC lines to generate patterned ventral forebrain organoids using a drug-treatment regimen that augments SHH signaling while concurrently inhibiting WNT signaling, as previously described (Bagley et al., 2017; Birey et al., 2017, 2022).

At the end of the patterning phase (day 24), brightfield microscopy revealed multiple germinal zone-like (rosettes) in organoids from both genotypes. At the two-month time point, ventral *ARHGEF6-KO* organoids were markedly smaller than isogenic controls, showing a significant reduction in area and density (**Fig. 4A–C**). In addition, *ARHGEF6-KO* organoids displayed a clear change in gross morphology, with an increased aspect ratio (a more elongated shape) and reduced roundness (less circular/compact profiles) compared with controls (**Fig. 4A, D, E**).

**Figure 4.**
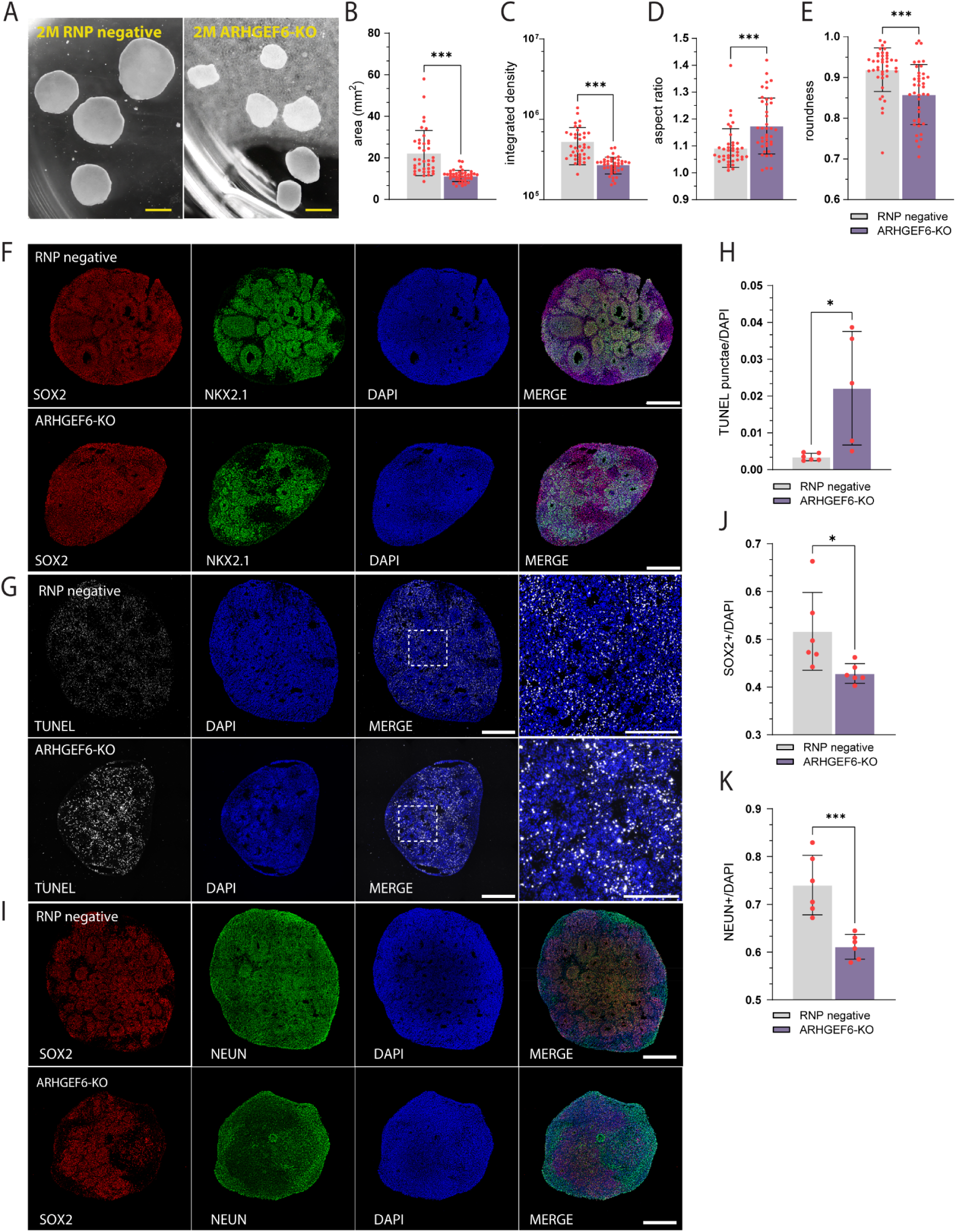
Characterization of ventral forebrain organoids from *ARHGEF6*-*KO* hiPSCs. **A**. Representative brightfield images of RNP negative (control) and *ARHGEF6-KO* human induced pluripotent stem cell (hiPSC)–derived organoids at 2 months (2M) of differentiation. Scale bars, 4 mm. **B–E**. Quantification of organoid size, density, and shape descriptors. 3 different batches of differentiation were analyzed. p-values = <0.0001 (area), <0.0001 (integrated density), <0.0001 (aspect ratio), <0.0001 (roundness). **F**. Immunofluorescence staining of 1-month ventral forebrain organoids showing expression of the neural progenitor marker SOX2 and the ventral telencephalic marker NKX2.1, confirming ventral forebrain identity. Nuclei are labeled with DAPI. Scale bars, 500 µm. **G**. Detection of apoptotic nuclei by TUNEL assay in RNP negative and *ARHGEF6-KO* 1-month organoids. Right panels show higher magnification of the boxed regions. Nuclei are labeled with DAPI. Scale bars, 500 µm. **H**. Average percentage of TUNEL punctae over the total number of nuclei. p-value = 0.015. At least 5 distinct sections from 5 different organoids per genotype from 3 different batches of differentiation were analyzed. **I**. Immunofluorescence staining for the neural progenitor marker SOX2 and the neuronal marker NEUN, indicating neuronal differentiation in RNP negative and *ARHGEF6-KO* 1-month organoids. Nuclei are labeled with DAPI. Scale bars, 500 µm. **J, K**. Quantification of SOX2⁺ progenitors and NEUN⁺ neurons normalized to the total number of nuclei in RNP negative and *ARHGEF6-KO* organoids. p-values = 0.028 (J), 0.0008 (K). At least 5 distinct sections from 6 different organoids per genotype from 3 different batches of differentiation were analyzed. p-values were calculated using unpaired t-test. * = p< 0.05, ** = p< 0.01, *** = p< 0.001. Data are presented as mean ± SEM. Each dot represents one organoid.

### 9. ARHGEF6 loss alters ventral forebrain organoid development

NKX2.1 expression is a hallmark of ventral telencephalic/MGE identity and is widely used to assess successful generation of MGE-like tissue, a principal source of cortical INs in humans (Hansen et al., 2013; Ma et al., 2013). Immunostaining for NKX2.1 in organoids from both genotypes confirmed successful ventral patterning (**Fig. 4F**).

To investigate cellular mechanisms that could contribute to the reduced size of *ARHGEF6-KO* organoids, we next assessed survival and cellular composition at one month. TUNEL staining revealed a significant increase in apoptotic cells in *ARHGEF6-KO* organoids (**Fig. 4G, H**). TUNEL punctae were distributed throughout the tissue and were particularly enriched in inter-rosette regions, suggesting that cell death may preferentially affect post-mitotic populations rather than proliferative progenitors within germinal zones. Consistent with this, quantification showed reduced numbers of both SOX2⁺ progenitors and NEUN⁺ neurons in *ARHGEF6-KO* organoids, with a more pronounced decrease in NEUN⁺ cells (**Fig. 4I-K**).

Collectively, these data indicate that ARHGEF6 loss is associated with a reduced NKX2.1⁺ progenitor pool and increased cell death, which may contribute to reduced neuronal output and could reflect impaired neurogenesis and/or decreased neuronal survival in ventral forebrain organoids.

### 10. *ARHGEF6-KO* INs exhibit impaired migration in dorsal-ventral assembloids

To assess the impact of ARHGEF6 loss on IN migration in a human context, we used dorsal–ventral assembloids to model ventral-to-dorsal migration. We transduced ventral organoids with lenti-DLX1/2b:eGFP to label INs and, at two months, fused them with dorsal organoids to generate dorsal–ventral assembloids, following established protocols (Bagley et al., 2017; Birey et al., 2017, 2022) (**Fig. 5A**). This system recapitulates migration of multiple MGE-derived IN subtypes and their characteristic saltatory behavior, consisting of alternating phases of rapid movement and pauses (Birey et al., 2017). To examine the behavior of migrating INs, we performed time-lapse recordings of dorsal-ventral assembloids cultured for 1-2 weeks after fusion (**Fig. 5B; Video S1, 2**). The eGFP^+^ ventral region was readily distinguishable from the dorsal region, allowing for clear visualization of the morphology of sparsely labeled migrating INs.

**Figure 5.**
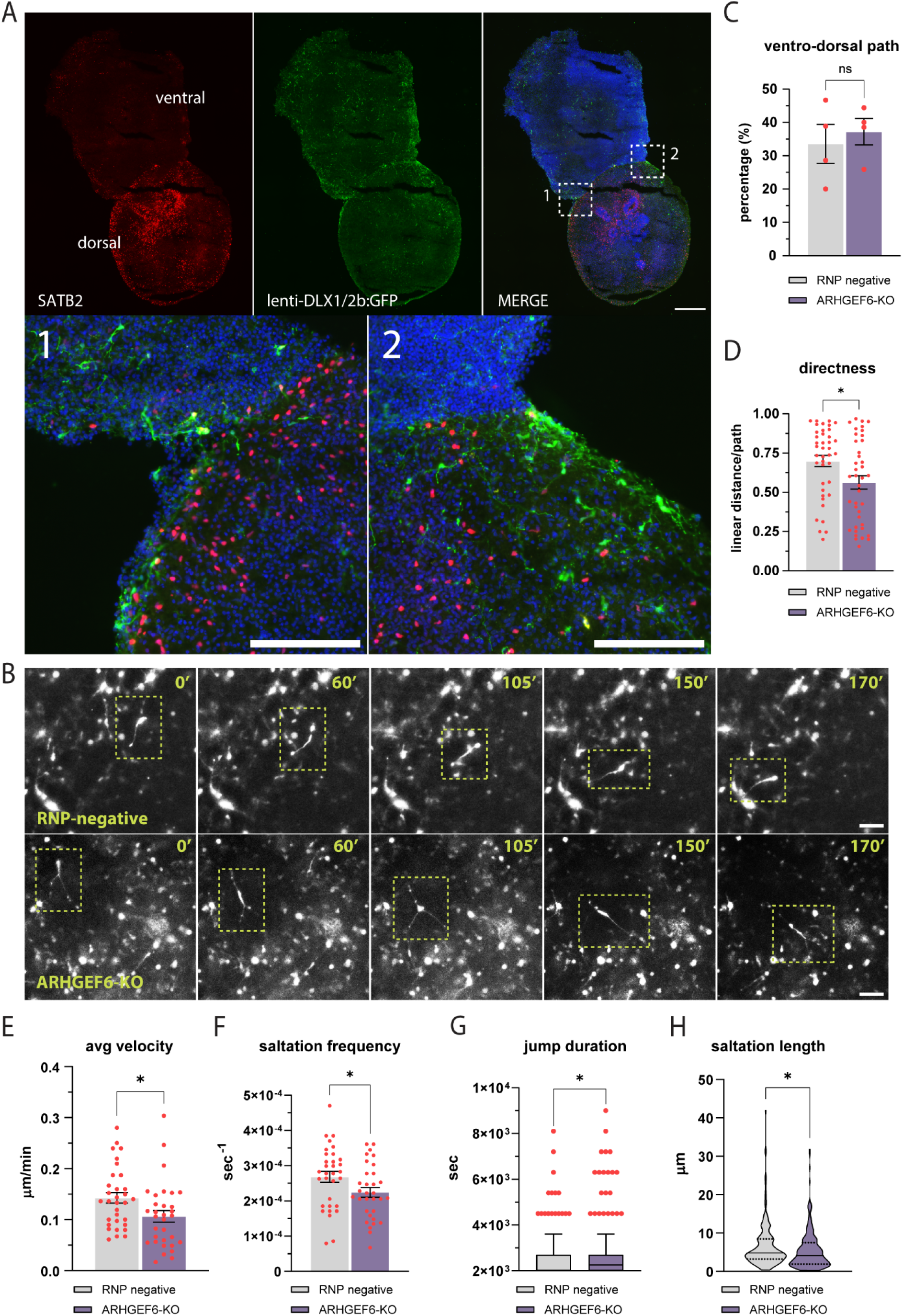
Impaired migration dynamics of DLX1/2b-GFP–labeled INs in ARHGEF6-deficient dorsal–ventral forebrain assembloids. **A**. Characterization of a fused dorsal–ventral assembloids by immunofluorescence. The dorsal compartment is labeled by SATB2 (red), whereas the ventral compartment expresses lenti-DLX1/2b-eGFP (green), marking ventral telencephalic IN progenitors. Nuclei are counterstained with DAPI (blue). Insets (1–2) show higher magnification of regions at the dorsal–ventral interface where GFP^+^ INs migrate from the ventral to the dorsal compartment. Scale bars, 500 µm, 150 (insets) µm. **B**. Representative frames from time-lapse imaging of migrating DLX1/2b-eGFP+ INs in fused assembloids, showing the saltatory migration behavior of individual INs in RNP negative (control) and *ARHGEF6-KO* assembloids. The tracked neuron is highlighted by dashed boxes in each frame. Scale bars, 100 µm. **C**. Quantification of the percentage of DLX1/2b-eGFP^+^ INs in the ventral compartment migrating toward the dorsal compartment. p-value = 0.624. **D–H**. Quantitative analysis of migration parameters of individual DLX1/2b-eGFP^+^ INs in RNP negative and *ARHGEF6-KO* assembloids. p-values = 0.016 (directness), 0.021 (average velocity), 0.038 (saltation frequency), 0.039 (jump (saltation event) duration), 0.043 (saltation length). At least 32 neurons from 3 independent assembloids per genotype from 2 different batches of differentiation were analyzed. p-values were calculated using unpaired t-test. * = p< 0.05, ** = p< 0.01, *** = p< 0.001. Data are presented as mean ± SEM. Each dot represents one tracked neuron. Violin plots show the distribution of values with median and quartiles indicated.

The fraction of DLX1/2b-eGFP⁺ INs migrating along the ventral-to-dorsal axis was comparable between genotypes (**Fig. 5C**). However, trajectory analysis revealed that *ARHGEF6-KO* INs migrated less directly, as reflected by reduced path directness (linear distance/path length) (**Fig. 5D**). Consistent with this, time-lapse imaging showed a reduction in overall average migration velocity (**Fig. 5E**) and altered saltatory dynamics (Birey et al., 2017), including fewer saltations, prolonged saltation events, and shorter saltation length (**Fig. 5F–H**).

Collectively, these results indicate that ARHGEF6 loss impairs the efficiency and coordination of IN migration in dorsal–ventral assembloids, despite preserved overall orientation.

### 11. *ARHGEF6-KO* INs show reduced branching

Given the morphological abnormalities observed in Arhgef6-deficient INs *in vivo*, we asked whether ARHGEF6 loss similarly affects neurite arborization in human INs. We therefore quantified neurite branching of DLX1/2b-eGFP⁺ INs from ventral organoids cultured for four months using Sholl analysis (**Fig. 6A)**. Compared with RNP negative controls, *ARHGEF6-KO* INs exhibited a reduced number of intersections in the proximal compartment (approximately 0–50 µm from the soma), whereas branching at larger distances was largely comparable between genotypes (**Fig. 6B**).

**Figure 6.**
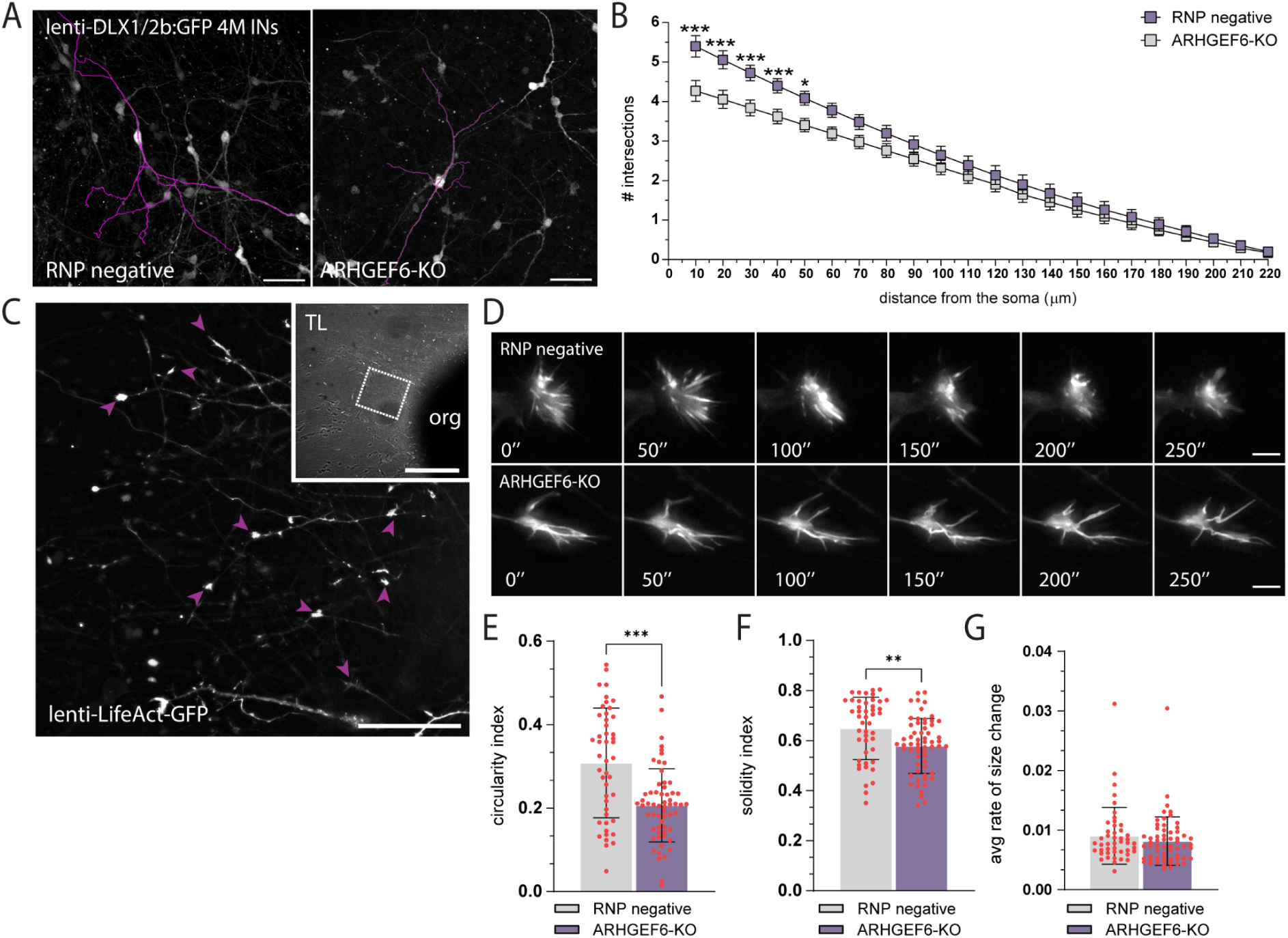
Impaired neuritogenesis and disrupted growth cone morphology in *ARHGEF6-KO* human neurons. **A**. Representative fluorescence images of reconstructed individual DLX1/2b-eGFP^+^ INs (purple skeletonization) from 4 months (4M) RNP negative (left) and *ARHGEF6-KO* (right) forebrain ventral organoids. Scale bar, 70 µm. **B**. Sholl analysis quantifying neurite length/complexity as the number of intersections across concentric radii (10–220 µm). p-values (from 10 to 220 µm) = 0.000003 (10 µm), 0.00003 (20 µm), 0.0003 (30 µm), 0.002 (40 µm), 0.008 (50 µm), 0.027 (60 µm), 0.070 (70 µm), 0.148 (80 µm), 0.263 (90 µm), 0.403 (100 µm), 0.548 (110 µm), 0.615 (120 µm), 0.569 (130 µm), 0.615 (140 µm), 0.679 (150 µm), 0.783 (160 µm), 0.814 (170 µm), 0.835 (180 µm), 0.848 (190 µm), 0.848 (200 µm), 0.848 (210 µm), 0.848 (220 µm). **C**. Representative image of neurites extending from a 1-month forebrain organoids transduced with LifeAct-GFP lentivirus. The inset contains a transmitted-light (TL) image of the organoid (org) plated showing migrating neurons and neuronal extensions. LifeAct-GFP^+^ growth cones are indicated by purple arrowheads. Scale bars, 50 µm, 200 µm (inset). **D**. Representative frames from time-lapse recordings of growth cones from RNP negative (top) and *ARHGEF6-KO* (bottom) LifeAct-GFP^+^ growth cones at 0’’, 50’’, 100’’, 150’’, 200’’, and 250’’. Scale bar, 5 µm. **E–G**. Quantification of growth cone shape descriptors and dynamics. p-values = < 0.001 (average circularity index), 0.003 (average solidity index), 0.299 (average rate of change in size). A total of 43 growth cones from 6 RNP negative organoids and 60 growth cones from 6 *ARHGEF6-KO* organoids, derived from 2 independent differentiation batches, were analyzed. Each dot represents one tracked growth cone. Data are shown as mean ± SEM. Statistical comparisons were performed using unpaired t-tests (E-G) and unpaired multiple t-tests corrected for False Discovery Rate (<1%) (B). *p < 0.05, **p < 0.01, ***p < 0.001.

The apparent discrepancy between the proximal branching defect observed in organoid-derived INs and the reduction of distal branching detected in primary neuronal cultures likely reflects differences in developmental maturation. Primary neurons have already undergone early stages of neurite specification *in vivo* prior to dissociation, revealing defects mainly in distal arborization. In contrast, organoid-derived neurons establish their neuritic arbor entirely *in vitro*, allowing earlier defects in proximal branching to become apparent. This result reinforces our previous findings, confirming that loss of ARHGEF6 reduces neurite complexity in both mouse and human models.

### 12. *ARHGEF6-KO* neurons exhibit altered growth cones morphology

Because migration and neurite branching depend on RAC1-driven cytoskeletal remodeling at the growth cone, we next asked whether ARHGEF6 loss impacts growth cone architecture in human neurons.

To visualize F-actin dynamics in live cells, we transduced one-month forebrain organoids with lenti–LifeAct-GFP and plated organoid-derived cells onto Geltrex-coated surfaces to promote adhesion and process extension. We then performed time-lapse imaging to monitor F-actin–rich growth cones in migrating neurons (**Fig. 6C, D; Video S3, 4**). Quantitative analysis showed that *ARHGEF6-KO* growth cones had reduced circularity and solidity (**Fig. 6E, F**), indicating a less compact morphology with a more elongated, protrusion-rich outline. In contrast, the average rate of size change over the imaging period was comparable between genotypes (**Fig. 6G**).

Collectively, these findings support a model in which ARHGEF6 contributes to cytoskeletal organization at the growth cone, providing a potential cellular basis for the migration and maturation deficits observed upon its loss.

## Discussion

In this study, we identify ARHGEF6 as a conserved regulator of forebrain GABAergic IN development in both mouse and human iPSC-derived systems, and show that loss of ARHGEF6 consistently impairs processes central to inhibitory circuit assembly, including migration, neurite branching, growth cone organization, electrophysiological maturation, and survival. These findings are particularly relevant given the established importance of GABAergic INs in the assembly and function of forebrain circuits. Although they represent only a minority of the total neuronal population, cortical and hippocampal INs regulate virtually all aspects of local circuit maturation and activity through their morphological, neurochemical, and functional diversity (Pelkey et al., 2017; Lim et al., 2018; Kirmse et al., 2022). Developmental disruption of IN number, positioning, subtype composition, or intrinsic properties can therefore produce long-lasting circuit-level consequences and has been implicated in multiple neurodevelopmental disorders (Marín, 2012; Yang et al., 2022; Marilovtseva et al., 2025).

The molecular function of ARHGEF6 previously described in literature is highly consistent with the phenotypes observed here. Cell migration, neuritogenesis, and growth cone navigation all require tight control of the actin cytoskeleton, a process largely governed by Rho-family GTPases and their regulators. In line with this, ARHGEF6-deficient INs showed abnormalities in multiple cytoskeleton-dependent processes, including reduced migration directness and speed, reduced neurite complexity, and abnormal growth cone morphology. These phenotypes were observed not only in the mouse model but also in dorsal–ventral assembloids, indicating that the function of ARHGEF6 in IN development is conserved across species. The concordance between mouse and human systems is particularly important given the longstanding uncertainty surrounding the relevance of ARHGEF6 dysfunction to human disease.

Our human assembloid experiments extend the mouse findings by showing that *ARHGEF6-KO* INs exhibit reduced migration efficiency and impaired saltatory behavior (Birey et al., 2017). This parallels our *in vivo* observations in embryonic mouse brain, where fewer INs entered the cortex and their leading processes showed reduced coherence along the tangential axis. Likewise, branching defects were observed in both species, although they were more distal in mouse primary neurons and more proximal in organoid-derived human INs. We interpret this apparent discrepancy as a difference in developmental context rather than a contradiction, as both systems converge on the same conclusion: loss of ARHGEF6 reduces IN neuritic complexity.

A further key finding of our study is the increase in developmental cell death in the absence of ARHGEF6. In the mouse, apoptosis was elevated in the GEs at E14.5 and in the hippocampus at E18.5; in human ventral organoids, TUNEL-positive cells were also increased, particularly outside rosette-like proliferative zones. At present, we cannot determine whether this survival defect is entirely cell-autonomous or also influenced by altered network activity. However, neuronal survival during development is shaped not only by an intrinsic basal rate of cell death but also, to a large extent, by activity-dependent mechanisms (Heck et al., 2008; Southwell et al., 2012; Wong et al., 2018; Causeret et al., 2018; Bitzenhofer et al., 2021), and INs themselves contribute critically to early cortical activity patterns, including spontaneous bursts and oscillations (Bonifazi et al., 2009; Isaacson & Scanziani, 2011; Blanquie et al., 2017). Because we observe altered migration, reduced IN numbers, and decreased firing in *Arhgef6* mutants, it is plausible that mutant INs fail to appropriately engage in immature activity patterns and are thereby rendered more vulnerable to developmental apoptosis.

This interpretation is reinforced by our electrophysiological data. In postnatal *Arhgef6* mutant mice, hippocampal INs showed a reduction in firing rate in response to current injection, as well as altered spike-frequency adaptation, overall consistent with a hypoexcitable phenotype. These changes occurred in the absence of major differences in passive membrane properties or single action potential waveform, suggesting a selective alteration in intrinsic firing behavior rather than a general loss of membrane integrity. Such a change is likely to affect local inhibitory function and could sum up with the synaptic plasticity defects already described in *Arhgef6*-deficient pyramidal neurons, including reduced LTP and enhanced LTD at the Schaffer collateral–CA1 synapse (Ramakers et al., 2012). Together, these findings support the idea that ARHGEF6 deficiency disrupts both excitatory and inhibitory components of the hippocampal circuitry, thereby increasing the likelihood of an excitation/inhibition imbalance relevant to cognitive dysfunction.

Our human organoid models also suggest that ARHGEF6 may act at earlier developmental stages, before overt IN migration begins. *ARHGEF6*-mutant ventral forebrain organoids were smaller, more elongated, less compact, and showed increased apoptosis along with reduced numbers of SOX2^+^ progenitors and NEUN^+^ neurons, raising the possibility of impaired progenitor maintenance and delayed neurogenesis. This is consistent with emerging evidence that Rho GTPases and their regulators contribute not only to migration and neurite dynamics, but also to neural progenitor proliferation, polarity, and fate decisions (Lian et al., 2019; Vidaki et al., 2012; Hass et al., 2025). More broadly, Rho GTPase signaling have been implicated in neurogenesis and microcephaly-associated phenotypes (Zuo et al., 2014; Jaffe and Hall, 2005; Reijnders et al., 2017; Chen et al., 2009; Duerinckx and Abramowicz, 2018). Although *ARHGEF6* mutations have not been clearly associated with microcephaly in patients, possibly due to the rarity of the condition, our organoid data suggest that ARHGEF6 may nonetheless influence early neurodevelopmental processes, including progenitor maintenance and neuronal output.

All these developmental findings are particularly relevant in light of the unresolved genetics of *ARHGEF6*-associated ID. *ARHGEF6* was initially proposed as the causal gene for nonsyndromic XLID46 (Kutsche et al., 2000; Yntema et al., 1998), but subsequent re-evaluation questioned this association because some reported variants are also present in apparently unaffected individuals and because direct functional evidence in human neurons has been limited (Piton et al., 2013). Clinical curation efforts have accordingly downgraded the evidence supporting a direct *ARHGEF6*–XLID46 relationship. However, the rarity of this condition makes it unlikely that human genetics alone will soon resolve the issue. In this context, our work provides precisely the kind of evidence that has been lacking: direct functional data in mouse and human inhibitory neurons showing that ARHGEF6 loss impairs core developmental processes highly relevant to cognition.

Finally, our results fit within a broader framework in which dysregulation of Rho GTPase signaling contributes to ID and related neurodevelopmental disorders. Variants in RAC1 itself cause a neurodevelopmental syndrome with ID, cortical malformations, and developmental delay, with both dominant-negative and gain-of-function alleles described (Banka et al., 2022; Haugh et al., 2021; Nishikawa et al., 2025; Priolo et al., 2023; Reijnders et al., 2017; Upadia et al., 2025). Together with the evidence on ARHGEF6 presented here, these studies reinforce the idea that perturbation of the RAC1 regulatory network has major developmental consequences in the human brain.

In conclusion, this study defines a developmental function for ARHGEF6 in forebrain GABAergic INs and provides mechanistic plausibility for how ARHGEF6 dysfunction could contribute to ID and related neurodevelopmental phenotypes. By demonstrating conserved defects in IN migration, maturation, cytoskeletal organization, and survival across mouse and human models, our work strengthens the link between disrupted inhibitory circuit assembly and cognitive impairment and provides experimental support for the relevance of ARHGEF6 dysfunction to human neurodevelopment.

## Materials and Methods

### Transcriptomic dataset acquisition and processing

The Allen Developing Human Brain Atlas: Developmental Transcriptome dataset, specifically the RNA-Seq Gencode v10 summarized to genes dataset, was downloaded from the Allen Institute for Brain Science (Allen Institute for Brain Science, 2010; available from brainspan.org; RRID:SCR_008083). Gene expression data were log-normalized using log_2_(RPKM + 1). A total of 30 samples from two unique donors were plotted in Figure 1A. The dataset from Wang et al. (2025), entitled Molecular and cellular dynamics of the developing human neocortex, was downloaded from CellxGene. Author-normalized gene expression values were used for this analysis; these values were calculated from raw snRNA-seq counts using SCTransform v2 (0.4.1) (Choudhary & Satija, 2022), as implemented in Seurat v4 (Hao et al., 2021; Wang et al., 2025). The dataset was filtered to include only postnatal data, resulting in the plotting of 114,216 of 232,328 total cells. Eighteen samples from nine unique subjects were plotted. Author-provided cell-type labels were used, and author-generated UMAP coordinates for each cell were obtained from the dataset. Please refer to the original publication for further details on data processing (Wang et al., 2025). Supplementary Table 1 contains postnatal *ARHGEF6* expression values for each cell type. The Mouse Whole Cortex and Hippocampus SMART-Seq dataset, together with the author-generated UMAP coordinates, was downloaded from the Allen Institute for Brain Science (Yao et al., 2021). This dataset includes scRNA-seq data from approximately 77,000 cells from male and female mice at around 8 weeks of age. Author-generated UMAP coordinates were used. Full names corresponding to acronyms were derived from the original publication and from the whole-mouse brain acronym list provided by the Allen Institute, available at brain-map.org. All analyses on this dataset were performed using the trimmed-means gene expression CSV file, in which the data were normalized as log_2_(CPM [exons + introns]) before calculation of mean expression for each cell-type cluster. Supplementary Table 2 contains *Arhgef6* expression values for each subclass. Multiple cell clusters were grouped into each subclass, and the trimmed mean expression of each cluster was averaged to obtain mean subclass-level expression. Please refer to the Allen Institute resource for additional information on data processing.

### Mouse strains

All animal procedures were approved by the local Animal Ethics Committee and the Ministry of Health. Animals were maintained in accordance with institutional animal welfare guidelines and legislation under veterinary supervision. The *Arhgef6-KO* mouse strain has been previously described (Ramakers et al., 2012) and was obtained from the INTRAFRONTIER EMMA Repository (EM:07499). Heterozygous and homozygous mutant mice are born at normal Mendelian frequency, appear grossly normal, are viable and fertile, mate at normal rates, and do not display evident neurological or motor impairments. Animals were maintained on a mixed C57BL/6 genetic background. The GAD67-eGFP reporter mouse strain has been previously described (DeDiego et al., 1994; Sakai & Miyazaki, 1997; Tamamaki et al., 2003). The progeny resulting from the cross between the reporter strain and the *Arhgef6-KO* strain were healthy and fertile.

### Brain preparation for histological analysis

For collection of postnatal brains, mice were anesthetized with Avertin (30 μl of pure Avertin in 400 μl PBS) and transcardially perfused with 10 ml PBS (pH 7.4), followed by 10 ml 4% (w/v) PFA in PBS (pH 7.4, adjusted with NaOH). Brains were dissected, post-fixed overnight at 4°C in 4% PFA, cryoprotected in 30% (w/v) sucrose in PBS followed by 15% (w/v) sucrose in PBS, embedded in OCT, and stored at −80°C until sectioning. OCT blocks were cut into 30 μm-thick coronal sections using a cryostat (Leica CM1950). Free-floating sections were collected in PBS in multiwell plates and stored at −20°C in cryoprotectant solution (30% [v/v] glycerol and 30% [v/v] ethylene glycol in 0.2 M phosphate buffer, pH 7.4) until processing. For collection of embryonic brains, embryos were obtained by cesarean section at E14.5 and E18.5 from anesthetized pregnant dams; detection of a vaginal plug was designated as E0.5. Embryos were transferred into PBS. Embryonic brains used for immunohistochemistry were dissected and fixed overnight at 4°C in 4% PFA, then cryoprotected overnight at 4°C in 30% (w/v) sucrose in PBS, followed by 15% (w/v) sucrose in PBS, embedded in OCT, and stored at −80°C until analysis. OCT blocks were cut into 20 μm-thick coronal sections and collected on adhesive glass slides.

### *In situ* hybridization

The murine *Arhgef6* antisense probe (Lein et al., 2007) was synthesized from adult mouse brain cDNA using the following primers (based on (Lein et al., 2007): forward 5′-CCTCGATTCTCCAGTAACCATC-3′ and reverse 5′-GGCCACTGATGAGTCCAACT-3′, with the reverse primer harboring the T7 promoter sequence 5′-GGTAATACGACTCACTATAGGG-3′ at the 5′ end. Probes were generated by two rounds of 35 PCR cycles using Phusion High-Fidelity DNA Polymerase (New England Biolabs, cat. M0530), followed by in vitro transcription with a digoxigenin (DIG) RNA Labeling Kit (Roche, cat. 11175025910). In situ hybridization was performed on 20 μm coronal cryosections from PFA-fixed wild-type mouse brains at the indicated time points using DIG-labeled antisense riboprobes (2 μg/ml), as previously described (Paganoni et al., 2022). Signal was revealed by colorimetric staining using 4-nitro blue tetrazolium chloride and 5-bromo-4-chloro-3-indolyl phosphate disodium salt (NBT/BCIP, Roche) in staining solution containing 100 mM NaCl, 50 mM MgCl2, 100 mM Tris-HCl (pH 9.8), 1% Tween 20, and 50% polyvinyl alcohol at 37°C.

### Primary cultures of hippocampal neurons

Two-well Ibidi slides were coated with 0.1 mg/ml poly-L-lysine (Sigma) in borate buffer (pH 8.5) and washed with deionized water. One day before establishing the culture, μ-slides were rinsed with MEM (Gibco) supplemented with 1% (v/v) sodium pyruvate 100× (Gibco), 20% (w/v) glucose, 1% (v/v) penicillin-streptomycin, and 10% (v/v) horse serum (Gibco). GAD67-eGFP and GAD67-eGFP;Arhgef6-KO pups at P0 were used to establish primary hippocampal neuronal cultures. Hippocampi were dissected from the brain after removal of the meninges under sterile conditions in a cold solution of 1% (v/v) HEPES in HBSS containing calcium and magnesium (Gibco). Hippocampi were washed in a cold solution containing 1% penicillin-streptomycin and 1% HEPES in HBSS with calcium and magnesium and incubated in 1 ml HBSS containing 25% Trypsin 0.25% (Gibco). Hippocampi were then washed twice in HBSS at 37°C for 10 min each. Tissue was dissociated by pipetting in a solution containing DNase (1:1,000; Promega). Cells were counted, and 430,000 cells were plated in each well containing Neurobasal medium (Gibco) supplemented with 1% penicillin-streptomycin, 2% (v/v) B27 (Gibco), and 0.25% (v/v) GlutaMAX (Gibco). Neurons were maintained at 37°C in a humidified atmosphere containing 5% CO2.

### Brain and primary cultures staining

Primary cortical cultures were fixed at 10 days *in vitro* (DIV10) with 4% PFA in PBS for 20 min at room temperature. Neurons were incubated for 1 h at room temperature in a blocking solution containing 5% goat serum and 0.1% Triton X-100 in PBS. The primary antibody (anti-GFP) was diluted in 3% goat serum and 0.1% Triton X-100 in PBS and incubated overnight at 4°C. Secondary antibodies were incubated for 1 h at room temperature. Finally, neurons were counterstained with DAPI before coverslips were mounted with Mowiol onto glass slides. Primary antibody: rabbit anti-GFP. Secondary antibody: Alexa Fluor 488 donkey anti-rabbit IgG (1:500; Invitrogen). Slides were examined with a Leica SP8 confocal microscope. Raw images were digitally processed in ImageJ (NIH, Bethesda, MD, USA) to normalize background, optimize contrast, rotate, and resize images. Morphological analysis of primary cultures was performed in ImageJ; neuronal arborization was quantified by Sholl analysis (Sholl, 1953) using the Sholl Analysis plugin (v1.0; Ghosh Lab Software). Terminal deoxynucleotidyl transferase dUTP nick end labeling (TUNEL; Click-iT™ TUNEL Alexa Fluor Imaging Assays for Microscopy & HCS, Invitrogen) was performed according to the manufacturer’s instructions on embryonic brain slices. Before imaging, embryonal sections were counterstained with DAPI and mounted with Mowiol on adhesive glass slides.Stained mouse brain sections were imaged using a 20× objective on a Leica SP8 confocal microscope and used for quantification on ImageJ.

### Whole-cell patch-clamp recording

For acute slice recordings, P90 mice were sacrificed by cervical dislocation. Brains were removed quickly and placed in an ice-cold sucrose-based cutting solution containing (in mM): 2.5 KCl, 1.25 NaH₂PO₄, 10 MgSO₄, 0.5 CaCl₂, 11 glucose, 234 sucrose, and 26 NaHCO₃. The solution was continuously bubbled with 95% O₂ / 5% CO₂ to maintain pH 7.3–7.4. Coronal somatosensory cortex slices (300 μm) were cut in ice-cold ACSF using a vibratome (Microm HM650V, Thermo Scientific) and then incubated for 30 min in ACSF containing 119 NaCl, 2.5 KCl, 26 NaHCO₃, 2.5 CaCl₂, 1.3 MgSO₄, 1.2 NaH₂PO₄, and 11 glucose (Liaci et al., 2022; Marcantoni et al., 2014). Slices recovered for 30–45min at 37 °C and then for an additional 1 h at room temperature before recordings. Patch electrodes were fabricated from borosilicate glass (Hilgenberg, Mansfield, Germany) and had a final resistance of 5–9 MΩ. For current-clamp recordings in both brain slices and primary cultured neurons, the internal solution contained 135 mM potassium gluconate, 5 mM NaCl, 2 mM MgCl_2_, 10 mM HEPES, 0.5 mM EGTA, 2 mM ATP-Tris, and 0.4 mM Tris-GTP. Whole-cell patch-clamp recordings from cortical INs in somatosensory cortex layers IV–VI were performed using an EPC-10 amplifier (HEKA Elektronik, Lambrecht, Germany). Traces were sampled at 10 kHz and low-pass filtered at 2 kHz with a Bessel filter. All experiments were performed at room temperature (22–24°C). Resting membrane potential (Vrest) and membrane capacitance (C_m_) were routinely measured upon establishment of whole-cell configuration. The membrane time constant (τ_m_) was calculated in Clampfit following a −30 pA current step injection. Cm was calculated according to C_m_ = τ_m_/R_in_. Action potential parameters were obtained by analyzing a series of spikes recorded during tonic firing lasting 1–2 min. Tonic firing was elicited by depolarizing the membrane with the minimum current required to reach the rheobase (Marcantoni et al., 2014). After reaching steady-state firing, at least five action potentials were selected and averaged for each cell. Action potential peak amplitude, half-width, maximum rising slope, and maximum repolarizing slope were measured using Clampfit software (Axon Instruments). Peak amplitude was measured from threshold to the action potential peak, and half-width was calculated at half-maximal height. To analyze the relationship between firing frequency and injected current, the membrane potential was adjusted to −70 mV and 20 current pulses of increasing intensity (from −30 to 160 pA; 500 ms duration) were injected. Mean firing frequency at each current step was calculated as the number of spikes per second. Rheobase was defined as the minimum current required to trigger one spike. Input resistance (R_in_) was calculated from the linear portion of the current-voltage relationship centered at the holding potential (−70 mV), using hyperpolarizing and depolarizing current steps from −30 to 30 pA in 10 pA increments.

### hiPSC mutagenesis

Human iPSCs DYS0100 (ATCC) were dissociated using TrypLE Select Enzyme (Gibco) and electroporated using a Lonza 4D-Nucleofector (program CM-113, solution P3) according to the manufacturer’s instructions (Umbach et al, 2022). Briefly, equal amounts of 100 μM crRNA and tracrRNA (Integrated DNA Technologies) were mixed and annealed to form gRNAs. A total of 150 pmol gRNA was complexed with 120 pmol Cas9 protein (Integrated DNA Technologies) to form ribonucleoprotein complexes (RNPs). The electroporation mixture was prepared as previously described (Ghetti et al., 2021). After electroporation, cells were plated onto Geltrex-coated plates and maintained in StemFlex medium supplemented with RevitaCell Supplement (Gibco) for 48 h. Monoclonal cell lines were established by single-cell sorting into 96-well plates followed by clonal expansion. Potential off-target sites for gRNA + 4 were analyzed using the Cas-OFFinder online algorithm with the following parameters: SpCas9 from Streptococcus pyogenes (5′-NGG-3′), mismatch number ≤ 4, DNA bulge size = 0, RNA bulge size = 0, and Homo sapiens (GRCh38/hg38) as the target genome. Genomic DNA was extracted using QuickExtract DNA Extraction Solution (Epicentre), and the target locus was amplified by PCR using Phusion High-Fidelity DNA Polymerase (Thermo Fisher). Oligonucleotides used to evaluate indels resulting from cleavage by one crRNA are listed below. Purified PCR products were sequenced and analyzed using Synthego ICE software (Brinkman et al., 2014, 2018; Conant et al., 2022; Kluesner et al., 2018). The following genomic regions were amplified and sequenced to validate CRISPR editing of ARHGEF6 and assess potential off-target effects. Primer sequences, amplicon size, and melting temperature (T_m_) are reported below. The target ARHGEF6 locus (chrX:+136747259) was amplified using the forward primer ATCACGAGAACAACTCGGC and the reverse primer GAGTGGGTCTGAGTATGCAC, generating a 480 bp amplicon with a T_m_ of 58°C. The first predicted off-target site (chrX:+106599275) was amplified using the forward primer GAAATGGTGGTTCCAGACAGC and the reverse primer GAACCGCACCACTCTGTTG, producing a 343 bp amplicon with a T_m_ of 59°C. The second predicted off-target site (chr6:-67012633) was amplified using the forward primer TCTGTAGCAAAACAACTGGC and the reverse primer TCACCTGCTAATGACCAAACAC, generating a 447 bp amplicon with a T_m_ of 58.55°C. The third predicted off-target site (chr4:+168553958) was amplified using the forward primer TGAACACCCATTGCATCCCCTC and the reverse primer AGGTGCCCAGAAGACCTTTATC, producing a 640 bp amplicon with a T_m_ of 59.16°C.

### hiPSC differentiation into neural progenitors and embryoid bodies

Human iPSCs (hiPSCs) were expanded on Geltrex-coated 6-well plates in Essential 8 (E8; Gibco) medium, which was changed daily. For passaging, cultures were washed with DPBS and exposed to 0.5 mM EDTA dissociation solution at 37°C until the cells and colonies appeared rounded but not fully detached. The dissociation buffer was then removed, colonies were collected and transferred to a 15 ml Falcon tube, centrifuged at 200 × g for 5 min, resuspended in E8 medium, and replated in 6-well plates. Cells were maintained at 37°C in 5% CO2, and medium was replaced daily. For embryoid body (EB) formation, hiPSCs were collected from one well of a 6-well plate at 70–80% confluence, centrifuged at 200 × g for 5 min, and the pellet was gently resuspended in complete E8 medium. Cells from each clone were plated in 4 ml complete E8 medium supplemented with 5 μM ROCK inhibitor Y-27632 into a 6-well plate pretreated with Pluronic acid. Plates were coated with 50 mg/ml Pluronic acid solution for 1 h at room temperature. The plate was gently tilted several times to distribute the cell suspension evenly, and cells were cultured under standard conditions (37°C, 5% CO2, 21% O2). EBs were checked daily, and medium was replaced every second day. During medium changes, EBs were collected into a 15 ml Falcon tube, allowed to settle by gravity, the supernatant was replaced, and the EBs were returned to the original wells. From day 2 to day 7, cultures were gradually shifted from E8 medium to Essential 6 (E6; Gibco) medium until EBs were maintained exclusively in E6. On day 7, EBs were transferred onto Geltrex-coated μ-Slide 2-well Ibidi chambers to allow adhesion and imaging analysis. E6 medium was replaced every other day. Neural induction, corresponding to the conversion of human iPSCs into neural progenitor cells (NPCs), was performed using PSC Neural Induction Medium (Gibco) according to the manufacturer’s instructions.

### hiPSC and hiPSC-derived cell staining

Cells plated in Geltrex-coated μ-Slide 2-well Ibidi chambers were fixed with 4% PFA for 30 min at room temperature and washed with PBS before incubation for 1 h in PBS containing 0.3% Triton X-100 (Sigma, T9284) and 6% BSA (Sigma, AA0281). Cells were incubated overnight at 4°C with primary antibodies, followed by incubation for 2 h at room temperature with secondary antibodies; both antibody solutions consisted of PBS containing 0.1% Triton X-100 and 2.5% BSA, with three washes before and after secondary antibody incubation. Primary antibodies used were mouse anti-OCT3/4 (1:1,000; Santa Cruz Biotechnology, sc-5279), rabbit anti-SOX2 (1:1,000; Abcam, AB97959), mouse anti-GATA4 (1:1,000; Santa Cruz Biotechnology, sc-25310), rabbit anti-TUJ1 (1:1,000; GeneTex, GTX130245), mouse anti-ACTA2 (1:1,000; Antibodies, A279072), mouse anti-NESTIN (1:1,000; R&D Systems, MAB1259), mouse anti-PAX6 (1:1,000; BD Pharmingen, 561462), and rabbit anti-FABP7 (Millipore, ABN14). For phalloidin staining, cells were washed in PBS and fixed in 4% formaldehyde in PBS for 30 min. Cells were permeabilized with 0.1% Triton X-100 in PBS for 5 min and blocked in 3% non-fat dry milk in PBS for 30 min. A total of 200 μl Phalloidin-FITC (1:500 in BSA 10× PBS) was added to each well for 90 min. Mowiol was used as mounting medium to preserve fluorescence. Coverslips were examined with a Leica SP8 laser scanning confocal microscope using 488 nm excitation. Images were analyzed in ImageJ software for corrected total cell fluorescence (CTCF), calculated from integrated density and background fluorescence measurements. Single rosettes were imaged for 14 h under bright-field illumination using a 20× objective on a Leica Thunder Microscope. Temperature and CO2 were maintained at 37°C and 5%, respectively, throughout the recording. Anisotropy was automatically calculated using the FibrilTool plugin in ImageJ (Boudaoud et al., 2014).

### Differentiation and assembly of dorsal and ventral organoids

Cortical and ventral differentiations from hiPSCs were performed as previously described (Birey et al., 2017; Sloan et al., 2018). For feeder-free cortical and ventral differentiation, hiPSCs were maintained on Geltrex-coated Costar plates in Essential 8 medium in a humidified atmosphere containing 5% CO2. Cells were passaged every 4–5 days using UltraPure 0.5 mM EDTA, pH 8.0 (Thermo Fisher Scientific, 15575020). On day 0, feeder-free human pluripotent stem cells at 80–90% confluence were dissociated into single cells with Accutase and reaggregated at 9,000 cells per well in ultra-low-attachment 96-well V-bottom plates (sBio PrimeSurface plate; Sumitomo Bakelite) in Essential 6 medium supplemented with the SMAD pathway inhibitors dorsomorphin (2.5 mM, Sigma-Aldrich, P5499) and SB431542 (10 mM, R&D Systems, 1614). Feeder-free cortical differentiation was performed as previously described (Yoon et al., 2019) using the protocol variant without XAV939. Feeder-free ventral differentiation was based on the feeder-free cortical protocol with the following additional treatments: XAV-939 (2.5 mM, Tocris, 3748) from day 3 to day 6; IWP-2 (2.5 mM, Selleck Chemicals, S7085) from day 7 to day 24; and SAG (100 nM, EMD Millipore, 566660) from day 13 to day 24. To promote progenitor differentiation, BDNF (20 ng/ml) and NT-3 (20 ng/ml) were added starting on day 25, with medium changes every other day. After day 43, medium was changed every 4–5 days using neural medium without growth factors. Assembly of cortical and ventral organoids to generate forebrain assembloids was performed as previously described (Birey et al., 2017; Sloan et al., 2018). 2-month-old ventral organoids were transduced with DLX1/2b-eGFP lentivirus (gift from S. Pasca and J. Rubenstein) as described in Paulsen et al. (2022). Bright-field images of all organoids were acquired from day 3 to day 60. ImageJ software was used to measure the area and perimeter of each organoid, and Prism software was used to plot the average size of organoids for each differentiation condition.

### Organoid IN morphological analysis

4-month-old ventral forebrain organoids transduced with DLX1/2b-eGFP lentivirus were imaged using a 25× water immersion objective on a Leica SP8 two-photon microscope. The SNT Neuroanatomy plugin (Arshadi et al., 2021) in ImageJ was used to perform Sholl analysis (Sholl, 1953).

### Organoid staining

Organoids were fixed with 4% PFA for 30 min at room temperature and then incubated overnight at 4°C in 30% sucrose solution. Organoids were embedded in Tissue-Tek O.C.T. compound (Sakura, 62550) and sectioned at 25 μm with a cryostat onto glass slides (Globe Scientific, 1354W). Slides were washed three times in PBS containing 0.1% Tween-20 (Sigma, P9416), then incubated for 1 h in PBS containing 0.3% Triton X-100 (Sigma, T9284) and 6% BSA (Sigma, AA0281). Slides were incubated overnight at 4°C with primary antibodies, followed by incubation for 2 h at room temperature with secondary antibodies. Both antibody solutions consisted of PBS containing 0.1% Triton X-100 and 2.5% BSA, with three washes before and after secondary antibody incubation. Slides were coverslipped using Fluoromount-G (EMS, 50-259-73). Primary antibodies used were goat anti-SOX2 (1:1,000; R&D Systems, AF2018), rabbit anti-NKX2.1 (1:500; Abcam, ab76013), rabbit anti-NEUN (1:200; Abcam, ab177487), mouse anti-SATB2 (1:100; Abcam, ab51502). TUNEL staining was performed according to the manufacturer’s instructions. Stained organoids were imaged using a 10× objective on a Leica Thunder Microscope and used for quantification.

### Organoids live imaging and analysis

Migration analysis. Live imaging of forebrain assembloids, whose ventral compartment had been previously transduced with LV-DLX1/2b-eGFP lentivirus, was performed using a 10× objective on a Leica THUNDER microscope, following the protocol described by Birey et al. (2017). All analyses were performed in ImageJ. Individual neurons were modeled as polylines defined by three points (A, B, and C), yielding two segments: AB, extending from the rear to the front of the soma, and BC, extending from the front of the soma to the tip of the leading process. A saltation event was defined as a sequence in which elongation of the total neuronal length, approximated as AB + BC, was followed by shortening.

Growth cone analysis. Approximately 1-month-old forebrain organoids were transduced with LifeAct-GFP lentivirus (Addgene, 51010), followed by a medium change 24 h after transduction. Five days after transduction, organoids were individually transferred to 1:70 Geltrex-coated μ-Slide 8-well glass-bottom plates (Ibidi, 80827). A volume of 1.5 μl Geltrex was applied directly to each organoid to promote attachment. To visualize growth cone dynamics in neurons extending processes outside the organoids, live imaging was performed over the following 10 days using a 63× oil immersion objective on a Leica Thunder Microscope. Time-lapse images were acquired in the GFP channel every 5 min for a total imaging period of 250 min for each selected field. Temperature and CO_2_ were maintained at 37°C and 5%, respectively, throughout the recording using a live-imaging chamber. Growth cones were manually selected for size and morphological analysis in ImageJ. The average rate of change in size was computed as the relative change between consecutive time points, (*S*_*t*+1_−*S*_*t*_)/*S*_*t*_, summed across the time series and normalized to the total imaging duration (250’’).

### Statistical analysis

Statistical analyses were performed using GraphPad Prism (GraphPad Software Inc.). The statistical test used for each experiment is indicated in the corresponding figure legends. Shapiro-Wilk and Kolmogorov-Smirnov tests were used to assess normality, and the F test was used to assess equality of variance; these results were used to select the appropriate statistical test. Data are presented as the mean ± standard error of the mean (SEM). In violin plots, the width of the violin reflects the distribution density of the data, the central line indicates the median, and the flanking lines indicate the interquartile range. Statistical significance was set at p < 0.05.

## Supporting information

Human postnatal ARHGEF6 expression by cell type.

Adult mouse Arhgef6 expression values by cell subclass.

Representative time-lapse imaging of an RNP negative LifeAct-GFP+ growth cone.

Representative time-lapse imaging of an ARHGEF6-KO LifeAct-GFP+ growth cone.

Representative time-lapse imaging of migrating RNP negative DLX1/2b-eGFP+ INs in fused assembloids.

Representative time-lapse imaging of migrating ARHGEF6-KO DLX1/2b-eGFP+ INs in fused assembloids.

## Acknowledgments

We thank Dr. Seth Ruffis (Optical Imaging Facility, Eli and Edythe Broad CIRM Center for Regenerative Medicine and Stem Cell Research, University of Southern California) and Dr. Marta Gai (Open Lab of Advanced Microscopy, OLMA@MBC, Molecular Biotechnology Center “Guido Tarone,” Università di Torino), and Federica Antico (Molecular Biotechnology Center “Guido Tarone,” Università di Torino) for their technical assistance. We thank Dr. S. Pasca and Dr. J. Rubenstein for generously providing the DLX1/2b-eGFP lentiviral vector.

## Data availability statement

The raw data supporting the conclusion of this article will be made available by the authors without undue reservation.

## Ethics statement

The animal study was reviewed and approved by the Italian Ministry of Health.

## Contributions

C.L., G.R.M., L.C., and G.Q. conceived the experiments. C.L., B.S., E.F., J.P.U., J.J.L., G.C., L.P., M.C., R.M., S.R., A.U., E.H., G.C., R.O., A.P., and V.T. performed the experiments. A.M., M.G., A.C., and L.C. provided materials and tools. C.L., G.R.M., and G.Q. supervised all aspects of the project. C.L., G.R.M., and G.Q. wrote the manuscript and designed the figures.

---

**Supplementary Figure 1.**
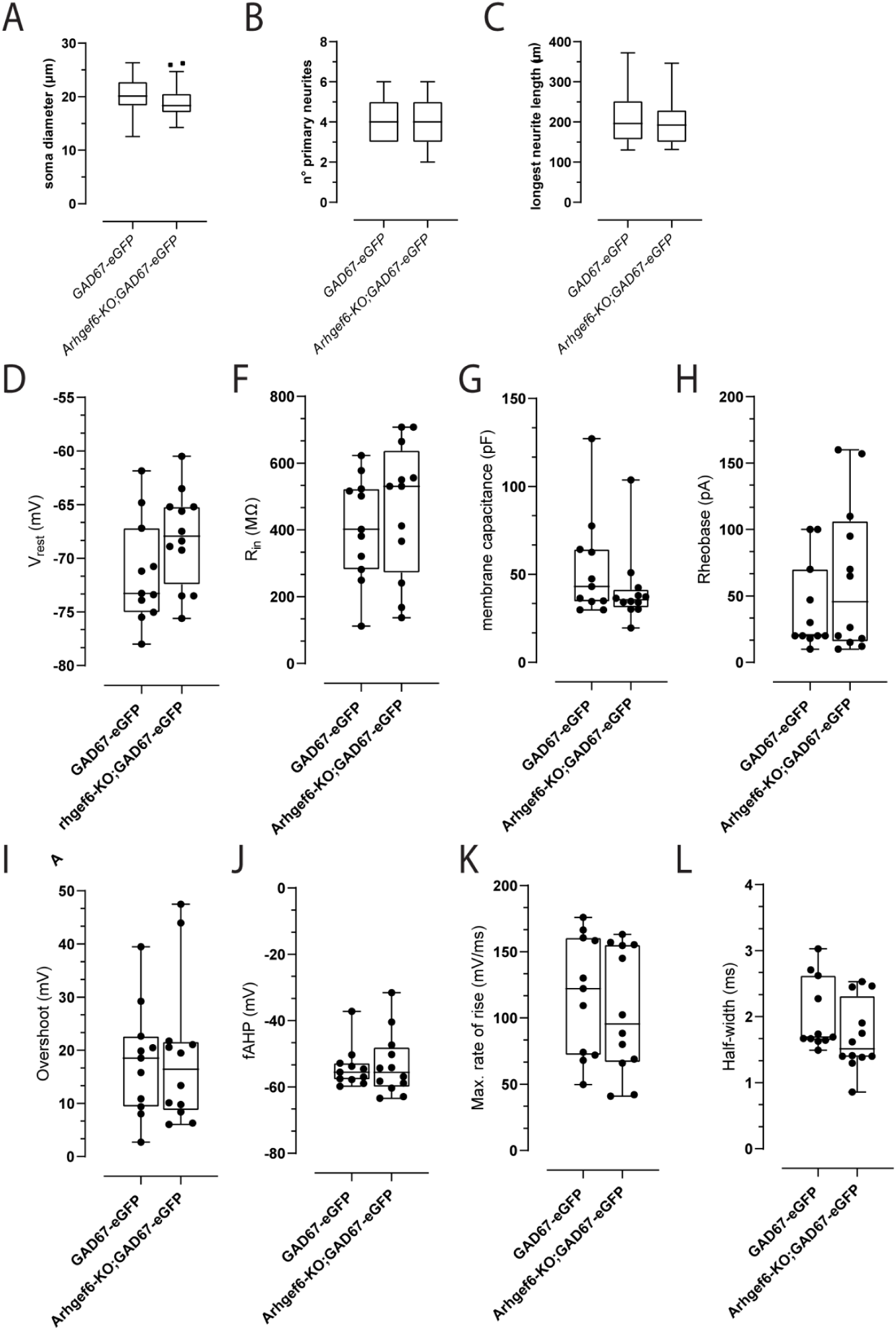
Morphological and intrinsic electrophysiological properties of *GAD67-eGFP* and *GAD67-eGFP;Arhgef6-KO* INs. **A–C**. Quantification of morphological parameters of reconstructed hippocampal primary INs after 10 days *in vitro* (DIV). p-values = 0.055 (soma diameter), 0.76 (number of primary neurites), 0.385 (longest neurite length). **D–H**. Passive membrane properties measured by whole-cell patch-clamp recordings in eGFP^+^ INs from control and *Arhgef6-KO* cultures. p-values = 0.105 (resting membrane potential, V_rest_), 0.464 (input resistance, R_in_), 0.287 (membrane capacitance), 0.275 (rheobase). **I–L**. Active membrane properties of recorded INs. p-values = 0.829 (action potential overshoot), 0.928 (after-hyperpolarization amplitude, AHP), 0.553 (maximum rate of rise of the action potential, Max. rate of rise), 0.182 (action potential half-width). p-values were calculated using unpaired t-test (F-I, K,L) and Mann-Whitney test (A-C, D, J). * = p< 0.05, ** = p< 0.01, *** = p< 0.001. Data are presented as mean ± SEM. Each dot represents an individual recorded neuron; box plots show median and interquartile range with whiskers indicating the data range.

**Supplementary Figure 2.**
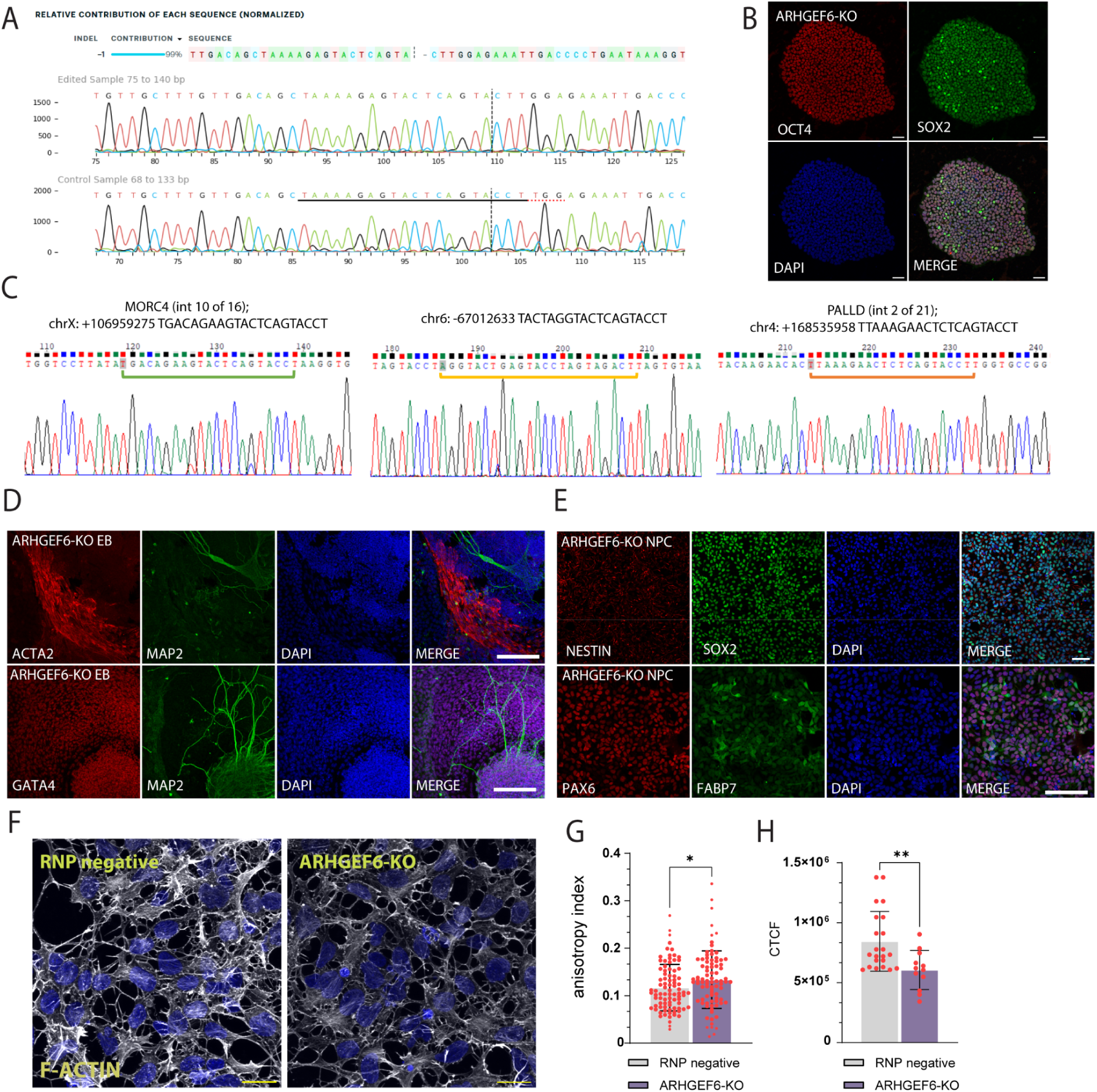
Mutagenesis of *ARHGEF6* in hiPSC and characterization of the line. **A**. Chromatogram of Sanger sequencing of human induced pluripotent stem cells (hiPSCs) ATCC-DYS0100, edited clone *ARHGEF6-KO* (top) and RNP negative isogenic control (bottom). The dashed line indicates the site of the CRISPR/Cas9 double-strand break. The mutation resulted in the deletion of a single cytosine, causing a frameshift mutation. **B**. Immunofluorescence staining for the expression of hiPSCs markers OCT4 (red) and SOX2 (green) in *ARHGEF6-KO* iPSCs. Nuclei were stained with DAPI (blue). Scale bars, 50 µm. **C**. Chromatograms from Sanger sequencing of the edited hiPSC clone showing no sequence changes at the top 3 predicted crRNA off-target sites. **D**. Immunofluorescence staining of marker genes for all three germ layers in embryoid bodies (EBs) obtained from *ARHGEF6-KO* hiPSCs. ACTA2 (mesoderm derivative, top, red), TUJ1/MAP2 (neuroectoderm, green), GATA4 (endoderm, bottom, red) and nuclei were stained with DAPI (blue). Scale bars, 200 µm. **E**. Immunofluorescence staining of neuroepithelial markers (top) Nestin (red), SOX2 (green), and radial glia markers (bottom) PAX6 (red) and FABP7 (green) in neural progenitor cells (NPCs) obtained from *ARHGEF6-KO* hiPSCs. Scale bars, 80 µm, 200 µm. **F**. Phalloidin-FITC staining of NPCs obtained from RNP negative (control) and *ARHGEF6*-KO hiPSC. Scale bars, 50 µm. **G**. Quantification of the anisotropy of F-Actin fibers in NPCs stained with Phalloidin-FITC. p-value = 0.0464. **H**. Quantification of polymerized F-actin by measuring the corrected total cell fluorescence (CTCF) of Phalloidin–FITC staining. p-value = 0.0056. p-values were calculated using unpaired t-test. * = p< 0.05, ** = p< 0.01, *** = p< 0.001. Data are presented as mean ± SEM. Each dot corresponds to a cell derived from 3 distinct batches of differentiation.

**Supplementary Table 1.** Human postnatal *ARHGEF6* expression by cell type. Single-nucleus RNA-sequencing (snRNA-seq) data from human postnatal donors (3 months - 13 years). Full snapshot of data used in Fig. 1B. Mean *ARHGEF6* expression per cell type across each postnatal timepoint and including all postnatal timepoints. Postnatal expression range: 0.0000 – 3.0445. 114,216 cells included from 18 samples across 9 unique subjects. 9,010 / 114,216 (7.9%) cells expressed *ARHGEF6* > 0. NaN occurs in the table when no cells of that cell type were present at the timepoint. EN-IT-Immature, intratelencephalic excitatory immature neurons; EN-L2_3-IT, layer 2/3 intratelencephalic excitatory neurons; EN-L4-IT, layer 4 intratelencephalic excitatory neurons; EN-L5-ET, layer 5 extratelencephalic excitatory neurons; EN-L5-IT, layer 5 intratelencephalic excitatory neurons; EN-L5_6-NP, layer 5/6 near-projecting excitatory neurons; EN-L6-CT, layer 6 corticothalamic excitatory neurons; EN-L6-IT, layer 6 intratelencephalic excitatory neurons; EN-L6b, layer 6b excitatory neurons; EN-Newborn, Newborn excitatory neurons; EN-Non-IT-Immature, immature non-intratelencephalic excitatory neuron;. IN-CGE-Immature, immature caudal ganglionic eminence-derived inhibitory neurons; IN-CGE-SNCG, immature caudal ganglionic eminence-derived gamma-synuclein inhibitory neurons; IN-CGE-VIP, caudal ganglionic eminence-derived vasoactive intestinal polypeptide inhibitory neurons; IN-MGE-Immature, immature medial ganglionic eminence-derived inhibitory neurons; IN-MGE-PV, medial ganglionic eminence-derived parvalbumin inhibitory neurons; IN-MGE-SST, medial ganglionic eminence-derived somatostatin inhibitory neurons; IN-Mix-LAMP5, mixed lysosomal-associated membrane protein family member 5 inhibitory neurons; IN-NCx_dGE-Immature, immature neocortex and dorsal ganglionic eminence-derived inhibitory neurons; IPC-EN, intermediate progenitor cell for excitatory neurons; OPC, oligodendrocyte precursor cells; RG-oRG, outer radial glial cells; RG-tRG, truncated radial glial cells; RG-vRG, ventricular radial glial cells; Tri-IPC, tripotential intermediate progenitor cells.

**Supplementary Table 2.** Adult mouse *Arhgef6* expression values by cell subclass. single-cell RNA-sequencing (scRNA-seq) data from ∼8 week old (∼P56) mice. Multiple cell clusters were grouped into each subclass, and the trimmed mean expression of each cluster was utilized to calculate subclass-level expression statistics. 73347 cells included. PV, parvalbumin; SST, somatostatin; SST CHOLD, somatostatin and chondrolectin; LAMP5, lysosomal-associated membrane protein family member 5; SNCG, gamma-synuclein; VIP, vasoactive intestinal polypeptide; MEIS2, meis homeobox 2; CR, cajal-retzius cell; L2 IT ENTl, layer 2 intratelencephalic lateral entorhinal area; L2 IT ENTm, layer 2 intratelencephalic medial entorhinal area; L2/3 IT CTX, layer 2/3 intratelencephalic isocortex; L2/3 IT ENTl, layer 2/3 intratelencephalic lateral entorhinal area; L2/3 IT PPP, layer 2/3 intratelencephalic postsubiculum-presubiculum-parasubiculum; L2/3 IT RHP, layer 2/3 intratelencephalic retrohippocampal region; L3 IT ENT, layer 3 intratelencephalic entorhinal area; L4 RSP-ACA, layer 4 retrosplenial area-anterior cingulate area; L4/5 IT CTX, layer 4/5 intratelencephalic isocortex; L5 IT CTX, layer 5 intratelencephalic isocortex; L5 PPP, layer 5 postsubiculum-presubiculum-parasubiculum; L5 PT CTX, layer 5 pyramidal tract isocortex; L5/6 IT TPE-ENT, layer 5/6 intratelencephalic temporal association areas-perirhinal area-ectorhinal area-entorhinal area; L5/6 NP CTX, layer 5/6 near-projecting isocortex; L6 CT CTX, layer 6 corticothalamic isocortex; L6 IT CTX, layer 6 intratelencephalic isocortex; L6 IT ENTl, layer 6 intratelencephalic lateral entorhinal area; L6b CTX, layer 6b isocortex; L6b/CT ENT, layer 6b corticothalamic entorhinal area; NP PPP, near-projecting postsubiculum-presubiculum-parasubiculum; NP SUB, near-projecting subiculum; CT SUB, corticothalamic subiculum; Car3, carbonic anhydrase 3; SUB-Pros, subiculum-prosubiculum; CA1-ProS, field cornu ammonis 1-prosubiculum; CA2-IG-FC, field cornu ammonis 2-fasciola cinerea-indusium griseum; CA3, field cornu ammonis 3; DG, dentate gyrus; Astro, astrocyte; Oligo, oligodendrocyte; Micro-PVM, microglia/perivascular macrophage; Endo, endothelial cell; SMC-Peri, smooth muscle cell perivascular area; VLMC, vascular leptomeningeal cell.

**Supplementary Video 1.** Representative time-lapse imaging of migrating RNP negative DLX1/2b-eGFP+ INs in fused assembloids.

**Supplementary Video 2.** Representative time-lapse imaging of migrating *ARHGEF6-KO* DLX1/2b-eGFP+ INs in fused assembloids.

**Supplementary Video 3.** Representative time-lapse imaging of an RNP negative LifeAct-GFP+ growth cone.

**Supplementary Video 4.** Representative time-lapse imaging of an *ARHGEF6-KO* LifeAct-GFP^+^ growth cone.

## References

Aerts, T., & Seuntjens, E. (2021). Novel perspectives on the development of the amygdala in rodents. Frontiers in Neuroanatomy, 15, 786679.

Arshadi, C., Günther, U., Eddison, M., Harrington, K. I. S., & Ferreira, T. A. (2021). SNT: A unifying toolbox for quantification of neuronal anatomy. Nature Methods, 18(4), 374–377.

Bagley, J. A., Reumann, D., Bian, S., Lévi-Strauss, J., & Knoblich, J. A. (2017). Fused cerebral organoids model interactions between brain regions. Nature Methods, 14(7), 743–751.

Banka, S., Bennington, A., Baker, M. J., Rijckmans, E., Clemente, G. D., Ansor, N. M., Sito, H., Prasad, P., Anyane-Yeboa, K., Badalato, L., Dimitrov, B., Fitzpatrick, D., Hurst, A. C. E., Jansen, A. C., Kelly, M. A., Krantz, I., Rieubland, C., Ross, M., Rudy, N. L., Sanz, J.,…Millard, T. H. (2022). Activating RAC1 variants in the switch II region cause a developmental syndrome and alter neuronal morphology. Brain, 145(12), 4232–4245.

Birey, F., Andersen, J., Makinson, C. D., Islam, S., Wei, W., Huber, N., Fan, H. C., Metzler, K. R. C., Panagiotakos, G., Thom, N., O’Rourke, N. A., Steinmetz, L. M., Bernstein, J. A., Hallmayer, J., Huguenard, J. R., & Paşca, S. P. (2017). Assembly of functionally integrated human forebrain spheroids. Nature, 545(7652), 54–59.

Birey, F., & Pașca, S. P. (2022). Imaging neuronal migration and network activity in human forebrain assembloids. STAR Protocols, 3(3), 101478.

Bitzenhofer, S. H., Pöpplau, J. A., Chini, M., Marquardt, A., & Hanganu-Opatz, I. L. (2021). A transient developmental increase in prefrontal activity alters network maturation and causes cognitive dysfunction in adult mice. Neuron, 109(8), 1350–1364.e6.

Blanquie, O., Kilb, W., Sinning, A., & Luhmann, H. J. (2017). Homeostatic interplay between electrical activity and neuronal apoptosis in the developing neocortex. Neuroscience, 358, 190–200.

Bonifazi, P., Goldin, M., Picardo, M. A., Jorquera, I., Cattani, A., Bianconi, G., Represa, A., Ben-Ari, Y., & Cossart, R. (2009). GABAergic hub neurons orchestrate synchrony in developing hippocampal networks. Science, 326(5958), 1419–1424.

Boudaoud, A., Burian, A., Borowska-Wykręt, D., Uyttewaal, M., Wrzalik, R., Kwiatkowska, D., & Hamant, O. (2014). FibrilTool, an ImageJ plug-in to quantify fibrillar structures in raw microscopy images. Nature Protocols, 9(2), 457–463.

Brinkman, E. K., Chen, T., Amendola, M., & van Steensel, B. (2014). Easy quantitative assessment of genome editing by sequence trace decomposition. Nucleic Acids Research, 42(22), e168.

Brinkman, E. K., Chen, T., de Haas, M., Holland, H. A., Akhtar, W., & van Steensel, B. (2018). Kinetics and fidelity of the repair of Cas9-induced double-strand DNA breaks. Molecular Cell, 70(5), 801–813.e6.

Cai, Y., Zhao, Z., Shi, M., Zheng, M., Gong, L., & He, M. (2024). Embryonic origins of forebrain oligodendrocytes revisited by combinatorial genetic fate mapping. eLife, 13, RP95406.

Causeret, F., Coppola, E., & Pierani, A. (2018). Cortical developmental death: Selected to survive or fated to die. Current Opinion in Neurobiology, 53, 35–42.

Chen, L., Liao, G., Waclaw, R. R., Burns, K. A., Linquist, D., Campbell, K., Zheng, Y., & Kuan, C. Y. (2007). Rac1 controls the formation of midline commissures and the competency of tangential migration in ventral telencephalic neurons. The Journal of neuroscience: the official journal of the Society for Neuroscience, 27(14), 3884–3893.

Chen, L., Melendez, J., Campbell, K., Kuan, C. Y., & Zheng, Y. (2009). Rac1 deficiency in the forebrain results in neural progenitor reduction and microcephaly. Developmental Biology, 325(1), 162–170.

Choudhary, S., & Satija, R. (2022). Comparison and evaluation of statistical error models for scRNA-seq. Genome Biology, 23(1), 27.

Conant, D., Hsiau, T., Rossi, N., Oki, J., Maures, T., Waite, K., Yang, J., Joshi, S., Kelso, R., Holden, K., Enzmann, B. L., & Stoner, R. (2022). Inference of CRISPR edits from Sanger trace data. The CRISPR Journal, 5(1), 123–130.

DeDiego, I., Smith-Fernández, A., & Fairén, A. (1994). Cortical cells that migrate beyond area boundaries: Characterization of an early neuronal population in the lower intermediate zone of prenatal rats. The European Journal of Neuroscience, 6(6), 983–997.

Duerinckx, S., & Abramowicz, M. (2018). The genetics of congenitally small brains. Seminars in Cell & Developmental Biology, 76, 76–85.

Eid, L., Lokmane, L., Raju, P. K., Tene Tadoum, S. B., Jiang, X., Toulouse, K., Lupien-Meilleur, A., Charron-Ligez, F., Toumi, A., Backer, S., Lachance, M., Lavertu-Jolin, M., Montseny, M., Lacaille, J. C., Bloch-Gallego, E., & Rossignol, E. (2025). Both GEF domains of the autism and developmental epileptic encephalopathy-associated Trio protein are required for proper tangential migration of GABAergic interneurons. Molecular Psychiatry, 30(4), 1338–1358.

Govek, E. E., Hatten, M. E., & Van Aelst, L. (2011). The role of Rho GTPase proteins in CNS neuronal migration. Developmental Neurobiology, 71(6), 528–553.

Ghetti, S., Burigotto, M., Mattivi, A., Magnani, G., Casini, A., Bianchi, A., Cereseto, A., & Fava, L. L. (2021). CRISPR/Cas9 ribonucleoprotein-mediated knockin generation in hTERT-RPE1 cells. STAR Protocols, 2(2), 100407.

Gupta, N. (2023). Deciphering intellectual disability. Indian Journal of Pediatrics, 90(2), 160–167.

Hansen, D. V., Lui, J. H., Flandin, P., Yoshikawa, K., Rubenstein, J. L., Alvarez-Buylla, A., & Kriegstein, A. R. (2013). Non-epithelial stem cells and cortical interneuron production in the human ganglionic eminences. Nature Neuroscience, 16(11), 1576–1587.

Hao, Y., Hao, S., Andersen-Nissen, E., Mauck, W. M., 3rd, Zheng, S., Butler, A., Lee, M. J., Wilk, A. J., Darby, C., Zager, M., Hoffman, P., Stoeckius, M., Papalexi, E., Mimitou, E. P., Jain, J., Srivastava, A., Stuart, T., Fleming, L. M., Yeung, B., Rogers, A. J.,…Satija, R. (2021). Integrated analysis of multimodal single-cell data. Cell, 184(13), 3573–3587.e29.

Hass, Y., Kniep, J., Hoffrichter, A., Marsoner, F., Eşiyok, N., Gasparotto, M., Xing, L., Loco Detto Gava, M. P., Artioli, A., Guida, C., Meuth, S. G., Huttner, W. B., Jabali, A., Heide, M., & Ladewig, J. (2025). ARHGAP11A maintains cortical progenitor identity through RHOA-ROCK signaling during human brain development. Cell Reports, 44(12), 116599.

Heck, N., Golbs, A., Riedemann, T., Sun, J. J., Lessmann, V., & Luhmann, H. J. (2008). Activity-dependent regulation of neuronal apoptosis in neonatal mouse cerebral cortex. Cerebral Cortex, 18(6), 1335–1349.

Haugh, I. M., Pineider, J. L., & Agim, N. G. (2021). Ichthyosiform changes in a patient with RAC1 mutation. Pediatric Dermatology, 38(6), 1590–1591.

Isaacson, J. S., & Scanziani, M. (2011). How inhibition shapes cortical activity. Neuron, 72(2), 231–243.

Jaffe, A. B., & Hall, A. (2005). Rho GTPases: Biochemistry and biology. Annual Review of Cell and Developmental Biology, 21, 247–269.

Keefe, F., Monzón-Sandoval, J., Rosser, A. E., Webber, C., & Li, M. (2023). Single-cell transcriptomics reveals conserved regulatory networks in human and mouse interneuron development. International Journal of Molecular Sciences, 24(9), 8122.

Kirmse, K., & Zhang, C. (2022). Principles of GABAergic signaling in developing cortical network dynamics. Cell Reports, 38(13), 110568.

Kluesner, M. G., Nedveck, D. A., Lahr, W. S., Garbe, J. R., Abrahante, J. E., Webber, B. R., & Moriarity, B. S. (2018). EditR: A method to quantify base editing from Sanger sequencing. The CRISPR Journal, 1(3), 239–250.

Kutsche, K., Yntema, H., Brandt, A., Jantke, I., Nothwang, H. G., Orth, U., Boavida, M. G., David, D., Chelly, J., Fryns, J. P., Moraine, C., Ropers, H. H., Hamel, B. C., van Bokhoven, H., & Gal, A. (2000). Mutations in ARHGEF6, encoding a guanine nucleotide exchange factor for Rho GTPases, in patients with X-linked mental retardation. Nature Genetics, 26(2), 247–250.

Lein, E. S., Hawrylycz, M. J., Ao, N., Ayres, M., Bensinger, A., Bernard, A., Boe, A. F., Boguski, M. S., Brockway, K. S., Byrnes, E. J., Chen, L., Chen, L., Chen, T. M., Chin, M. C., Chong, J., Crook, B. E., Czaplinska, A., Dang, C. N., Datta, S., Dee, N. R.,…Jones, A. R. (2007). Genome-wide atlas of gene expression in the adult mouse brain. Nature, 445(7124), 168–176.

Liaci, C., Camera, M., Caslini, G., Rando, S., Contino, S., Romano, V., & Merlo, G. R. (2021). Neuronal cytoskeleton in intellectual disability: From systems biology and modeling to therapeutic opportunities. International Journal of Molecular Sciences, 22(11), 6167.

Liaci, C., Camera, M., Zamboni, V., Sarò, G., Ammoni, A., Parmigiani, E., Ponzoni, L., Hidisoglu, E., Chiantia, G., Marcantoni, A., Giustetto, M., Tomagra, G., Carabelli, V., Torelli, F., Sala, M., Yanagawa, Y., Obata, K., Hirsch, E., & Merlo, G. R. (2022). Loss of ARHGAP15 affects the directional control of migrating interneurons in the embryonic cortex and increases susceptibility to epilepsy. Frontiers in Cell and Developmental Biology, 10, 875468.

Lian, G., Wong, T., Lu, J., Hu, J., Zhang, J., & Sheen, V. (2019). Cytoskeletal associated Filamin A and RhoA affect neural progenitor specification during mitosis. Cerebral Cortex, 29(3), 1280–1290.

Lim, L., Mi, D., Llorca, A., & Marín, O. (2018). Development and functional diversification of cortical interneurons. Neuron, 100(2), 294–313.

Ma, T., Wang, C., Wang, L., Zhou, X., Tian, M., Zhang, Q., Zhang, Y., Li, J., Liu, Z., Cai, Y., Liu, F., You, Y., Chen, C., Campbell, K., Song, H., Ma, L., Rubenstein, J. L., & Yang, Z. (2013). Subcortical origins of human and monkey neocortical interneurons. Nature Neuroscience, 16(11), 1588–1597.

Maia, N., Nabais Sá, M. J., Melo-Pires, M., de Brouwer, A. P. M., & Jorge, P. (2021). Intellectual disability genomics: Current state, pitfalls and future challenges. BMC Genomics, 22(1), 909.

Manser, E., Loo, T. H., Koh, C. G., Zhao, Z. S., Chen, X. Q., Tan, L., Tan, I., Leung, T., & Lim, L. (1998). PAK kinases are directly coupled to the PIX family of nucleotide exchange factors. Molecular Cell, 1(2), 183–192.

Marcantoni, A., Raymond, E. F., Carbone, E., & Marie, H. (2014). Firing properties of entorhinal cortex neurons and early alterations in an Alzheimer’s disease transgenic model. Pflügers Archiv: European Journal of Physiology, 466(7), 1437–1450.

Marilovtseva, E. V., Abdurazakov, A., Kurishev, A. O., Mikhailova, V. A., & Golimbet, V. E. (2025). The role of GABA pathway components in pathogenesis of neurodevelopmental disorders. International Journal of Molecular Sciences, 26(19), 9492.

Marín, O. (2012). Interneuron dysfunction in psychiatric disorders. Nature Reviews Neuroscience, 13(2), 107–120.

Marín, O. (2013). Cellular and molecular mechanisms controlling the migration of neocortical interneurons. The European journal of neuroscience, 38(1), 2019–2029.

Marín, O., & Rubenstein, J. L. (2001). A long, remarkable journey: tangential migration in the telencephalon. Nature reviews. Neuroscience, 2(11), 780–790.

Martino, A., Ettorre, M., Musilli, M., Lorenzetto, E., Buffelli, M., & Diana, G. (2013). Rho GTPase-dependent plasticity of dendritic spines in the adult brain. Frontiers in Cellular Neuroscience, 7, 62.

Meseke, M., Rosenberger, G., & Förster, E. (2013). Reelin and the Cdc42/Rac1 guanine nucleotide exchange factor αPIX/Arhgef6 promote dendritic Golgi translocation in hippocampal neurons. The European Journal of Neuroscience, 37(9), 1404–1412.

Meyer, M. A. (2014). Highly expressed genes within hippocampal sector CA1: Implications for the physiology of memory. Neurology International, 6(2), 5388.

Nishikawa, M., Hayashi, S., Nakayama, A., Nishio, Y., Shiraki, A., Ito, H., Maruyama, K., Muramatsu, Y., Ogi, T., Mizuno, S., & Nagata, K. I. (2025). Pathophysiological significance of the p.E31G variant in RAC1 responsible for a neurodevelopmental disorder with microcephaly. Biochimica et Biophysica Acta: Molecular Basis of Disease, 1871(1), 167520.

Nodé-Langlois, R., Muller, D., & Boda, B. (2006). Sequential implication of the mental retardation proteins ARHGEF6 and PAK3 in spine morphogenesis. Journal of Cell Science, 119(Pt 23), 4986–4993.

Paganoni, A. J. J., Amoruso, F., Porta Pelayo, J., Calleja-Pérez, B., Vezzoli, V., Duminuco, P., Caramello, A., Oleari, R., Fernández-Jaén, A., & Cariboni, A. (2022). A novel loss-of-function SEMA3E mutation in a patient with severe intellectual disability and cognitive regression. International Journal of Molecular Sciences, 23(10), 5632.

Paredes, M. F., James, D., Gil-Perotin, S., Kim, H., Cotter, J. A., Ng, C., Sandoval, K., Rowitch, D. H., Xu, D., McQuillen, P. S., Garcia-Verdugo, J. M., Huang, E. J., & Alvarez-Buylla, A. (2016). Extensive migration of young neurons into the infant human frontal lobe. Science, 354(6308), aaf7073.

Paulsen, B., Velasco, S., Kedaigle, A. J., Pigoni, M., Quadrato, G., Deo, A. J., Adiconis, X., Uzquiano, A., Sartore, R., Yang, S. M., Simmons, S. K., Symvoulidis, P., Kim, K., Tsafou, K., Podury, A., Abbate, C., Tucewicz, A., Smith, S. N., Albanese, A., Barrett, L.,…Arlotta, P. (2022). Autism genes converge on asynchronous development of shared neuron classes. Nature, 602(7896), 268–273.

Pelkey, K. A., Chittajallu, R., Craig, M. T., Tricoire, L., Wester, J. C., & McBain, C. J. (2017). Hippocampal GABAergic inhibitory interneurons. Physiological Reviews, 97(4), 1619–1747.

Piton, A., Redin, C., & Mandel, J. L. (2013). XLID-causing mutations and associated genes challenged in light of data from large-scale human exome sequencing. American Journal of Human Genetics, 93(2), 368–383.

Prakash, N., Abu Irqeba, A., & Corbin, J. G. (2025). Development and function of the medial amygdala. Trends in Neurosciences, 48(1), 22–32.

Priolo, M., Zara, E., Radio, F. C., Ciolfi, A., Spadaro, F., Bellacchio, E., Mancini, C., Pantaleoni, F., Cordeddu, V., Chiriatti, L., Niceta, M., Africa, E., Mammì, C., Melis, D., Coppola, S., & Tartaglia, M. (2023). Clinical profiling of MRD48 and functional characterization of two novel pathogenic RAC1 variants. European Journal of Human Genetics, 31(7), 805–814.

Ramakers, G. J., Wolfer, D., Rosenberger, G., Kuchenbecker, K., Kreienkamp, H. J., Prange-Kiel, J., Rune, G., Richter, K., Langnaese, K., Masneuf, S., Bösl, M. R., Fischer, K. D., Krugers, H. J., Lipp, H. P., van Galen, E., & Kutsche, K. (2012). Dysregulation of Rho GTPases in the αPix/Arhgef6 mouse model of X-linked intellectual disability is paralleled by impaired structural and synaptic plasticity and cognitive deficits. Human Molecular Genetics, 21(2), 268–286.

Reijnders, M. R. F., Ansor, N. M., Kousi, M., Yue, W. W., Tan, P. L., Clarkson, K., Clayton-Smith, J., Corning, K., Jones, J. R., Lam, W. W. K., Mancini, G. M. S., Marcelis, C., Mohammed, S., Pfundt, R., Roifman, M., Cohn, R., Chitayat, D., Deciphering Developmental Disorders Study, Millard, T. H., Katsanis, N.,…Banka, S. (2017). RAC1 missense mutations in developmental disorders with diverse phenotypes. American Journal of Human Genetics, 101(3), 466–477.

Sakai, K., & Miyazaki, J.-i. (1997). A transgenic mouse line that retains Cre recombinase activity in mature oocytes irrespective of the cre transgene transmission. Biochemical and Biophysical Research Communications, 237(2), 318–324.

Sholl, D. A. (1953). Dendritic organization in the neurons of the visual and motor cortices of the cat. Journal of Anatomy, 87(4), 387–406.

Sloan, S. A., Andersen, J., Pașca, A. M., Birey, F., & Pașca, S. P. (2018). Generation and assembly of human brain region-specific three-dimensional cultures. Nature Protocols, 13(9), 2062–2085.

Southwell, D. G., Paredes, M. F., Galvao, R. P., Jones, D. L., Froemke, R. C., Sebe, J. Y., Alfaro-Cervello, C., Tang, Y., Garcia-Verdugo, J. M., Rubenstein, J. L., Baraban, S. C., & Alvarez-Buylla, A. (2012). Intrinsically determined cell death of developing cortical interneurons. Nature, 491(7422), 109–113.

Sun, X., Wang, L., Wei, C., Sun, M., Li, Q., Meng, H., Yue, W., Zhang, D., & Li, J. (2021). Dysfunction of Trio GEF1 involves in excitatory/inhibitory imbalance and autism-like behaviors through regulation of interneuron migration. Molecular Psychiatry, 26(12), 7621–7640.

Tamamaki, N., Yanagawa, Y., Tomioka, R., Miyazaki, J., Obata, K., & Kaneko, T. (2003). Green fluorescent protein expression and colocalization with calretinin, parvalbumin, and somatostatin in the GAD67-GFP knock-in mouse. The Journal of Comparative Neurology, 467(1), 60–79.

Tejada-Simon, M. V. (2015). Modulation of actin dynamics by Rac1 to target cognitive function. Journal of Neurochemistry, 133(6), 767–779.

Tivodar, S., Kalemaki, K., Kounoupa, Z., Vidaki, M., Theodorakis, K., Denaxa, M., Kessaris, N., de Curtis, I., Pachnis, V., & Karagogeos, D. (2015). Rac-GTPases Regulate Microtubule Stability and Axon Growth of Cortical GABAergic Interneurons. Cerebral cortex (New York, N.Y.: 1991), 25(9), 2370–2382.

Uhlén, M., Fagerberg, L., Hallström, B. M., Lindskog, C., Oksvold, P., Mardinoglu, A., Sivertsson, Å., Kampf, C., Sjöstedt, E., Asplund, A., Olsson, I., Edlund, K., Lundberg, E., Navani, S., Szigyarto, C. A., Odeberg, J., Djureinovic, D., Takanen, J. O., Hober, S., Alm, T.,…Pontén, F. (2015). Proteomics. Tissue-based map of the human proteome. Science, 347(6220), 1260419.

Uhlen, M., Zhang, C., Lee, S., Sjöstedt, E., Fagerberg, L., Bidkhori, G., Benfeitas, R., Arif, M., Liu, Z., Edfors, F., Sanli, K., von Feilitzen, K., Oksvold, P., Lundberg, E., Hober, S., Nilsson, P., Mattsson, J., Schwenk, J. M., Brunnström, H., Glimelius, B.,…Ponten, F. (2017). A pathology atlas of the human cancer transcriptome. Science, 357(6352), eaan2507.

Umbach, A., Maule, G., Kheir, E., Cutarelli, A., Foglia, M., Guarrera, L., Fava, L. L., Conti, L., Garattini, E., Terao, M., & Cereseto, A. (2022). Generation of corrected hiPSC clones from a Cornelia de Lange syndrome (CdLS) patient through CRISPR-Cas-based technology. Stem Cell Research & Therapy, 13(1), 440.

Upadia, J., Liu, J., Bier, C., Chenevert, M., & Li, Y. (2025). Diverse clinical presentation of RAC1-related intellectual developmental disorder. American Journal of Medical Genetics Part A, 197(5), e63991.

Vidaki, M., Tivodar, S., Doulgeraki, K., Tybulewicz, V., Kessaris, N., Pachnis, V., & Karagogeos, D. (2012). Rac1-dependent cell cycle exit of MGE precursors and GABAergic interneuron migration to the cortex. Cerebral Cortex, 22(3), 680–692.

Wang, L., Wang, C., Moriano, J. A., Chen, S., Zuo, G., Cebrián-Silla, A., Zhang, S., Mukhtar, T., Wang, S., Song, M., de Oliveira, L. G., Bi, Q., Augustin, J. J., Ge, X., Paredes, M. F., Huang, E. J., Alvarez-Buylla, A., Duan, X., Li, J., & Kriegstein, A. R. (2025). Molecular and cellular dynamics of the developing human neocortex. Nature, 647(8088), 169–178.

Wonders, C. P., & Anderson, S. A. (2006). The origin and specification of cortical interneurons. Nature reviews. Neuroscience, 7(9), 687–696.

Wong, F. K., Bercsenyi, K., Sreenivasan, V., Portalés, A., Fernández-Otero, M., & Marín, O. (2018). Pyramidal cell regulation of interneuron survival sculpts cortical networks. Nature, 557(7707), 668–673.

Yang, J., Yang, X., & Tang, K. (2022). Interneuron development and dysfunction. The FEBS Journal, 289(8), 2318–2336.

Yao, Z., van Velthoven, C. T. J., Nguyen, T. N., Goldy, J., Sedeno-Cortes, A. E., Baftizadeh, F., Bertagnolli, D., Casper, T., Chiang, M., Crichton, K., Ding, S. L., Fong, O., Garren, E., Glandon, A., Gouwens, N. W., Gray, J., Graybuck, L. T., Hawrylycz, M. J., Hirschstein, D., Kroll, M.,…Zeng, H. (2021). A taxonomy of transcriptomic cell types across the isocortex and hippocampal formation. Cell, 184(12), 3222–3241.e26.

Yntema, H. G., Hamel, B. C., Smits, A. P., van Roosmalen, T., van den Helm, B., Kremer, H., Ropers, H. H., Smeets, D. F., & van Bokhoven, H. (1998). Localisation of a gene for non-specific X linked mental retardation (MRX46) to Xq25-q26. Journal of Medical Genetics, 35(10), 801–805.

Yoon, S. J., Elahi, L. S., Pașca, A. M., Marton, R. M., Gordon, A., Revah, O., Miura, Y., Walczak, E. M., Holdgate, G. M., Fan, H. C., Huguenard, J. R., Geschwind, D. H., & Pașca, S. P. (2019). Reliability of human cortical organoid generation. Nature Methods, 16(1), 75–78.

Zhou, W., Li, X., & Premont, R. T. (2016). Expanding functions of GIT Arf GTPase-activating proteins, PIX Rho guanine nucleotide exchange factors and GIT-PIX complexes. Journal of Cell Science, 129(10), 1963–1974.

Zuo, Y., Oh, W., & Frost, J. A. (2014). Controlling the switches: Rho GTPase regulation during animal cell mitosis. Cellular Signalling, 26(12), 2998–3006.

